# Adolescent development accelerates responses to input in human neocortical neurons

**DOI:** 10.64898/2026.09.16.746058

**Authors:** Rennie M Kendrick, Jason K Clark, Emmalyn P Leonard, Nivetha Ramasamy, Sabrina S DeLope, H. Westley Phillips, Vivek P Buch, Kelly B Mahaney, Gerald A Grant, Laura M Prolo, Scott F Owen

## Abstract

Through childhood and adolescence, profound changes to the physiology of individual neurons accompany large-scale network changes in the mammalian neocortex. These physiological changes are well understood in rodent models but far less is known in the human neocortex. Here we combine patch-clamp electrophysiology and single-cell sequencing (Patch-Seq) in neurosurgically-resected pediatric human brain slices and age-matched mouse brain slices to elucidate the unique developmental trajectory of human neurons. We find that human Layer 2/3 pyramidal neurons show distinctive postnatal changes in neuronal physiology that align with the more directed, feedforward network architecture of human neocortex relative to the mouse. Human-specific changes to spike train dynamics include faster spike latencies and selective acceleration of early spiking. By applying linear modeling to our Patch-Seq data, we identify genes that predict physiological variation across single cells. This unbiased approach unexpectedly identifies BK-type calcium-activated potassium channels as key drivers of human postnatal changes in spike train dynamics between childhood and adolescence. We further test this pathway through pharmacology and computational modeling. Together, our results reveal novel mechanisms of postnatal maturation in human neocortical neurons and demonstrate a new application of Patch-Seq to uncover gene-physiology relationships at single-cell resolution.

## Main

Across early childhood and adolescence, a coordinated set of molecular, cellular, and network transformations fundamentally restructure mammalian neocortical function.^1–19^ Changes to the intrinsic physiology of neurons are an established element of postnatal development in rodents,^8,9^ while defects in this maturation can lead to neurodevelopmental disorders such as autism,^20,21^ obsessive-compulsive disorder,^22–24^ and schizophrenia.^21,25^ Despite insights from rodent models,^8,9^ we still know very little about the postnatal changes in neuronal physiology in the juvenile human neocortex or their underlying mechanisms.^17,26–28^ This creates a barrier to translation of treatments for neurodevelopmental disorders to humans.

Here we combined whole-cell, brain slice patch-clamp recordings with single cell sequencing (Patch-Seq^29–31^) or pharmacology in *ex vivo* pediatric human and mouse neocortex to interrogate postnatal changes in physiology and underlying molecular mechanisms. This marks the first pediatric human Patch-Seq dataset and the first cross-species pediatric physiology dataset to our knowledge. We focused our experiments on Layer 2/3 (L2/3) pyramidal neurons because of their pronounced evolutionary divergence from rodent models in adult L2/3, including increased molecular diversity and distinctive physiology.^15,32–42^ More broadly, adult human L2/3 shows pronounced physical expansion^15,43,44^ and unique network architecture^38,39,45^ relative to mouse.

We uncovered human-specific changes in spike train dynamics that accelerate response speeds and early spike rates across the postnatal period. These physiological changes align with the more rapid, feedforward network architecture observed in adult human L2/3 relative to mouse.^39^ Linear modeling on our Patch-Seq dataset identified molecular candidates that could drive these human-specific developmental changes,^29,46–48^ including the BK-type calcium-activated potassium channel. Complementary pharmacology and *in silico* modeling provided causal evidence supporting a BK-mediated mechanism for these human-specific developmental changes.^49^

## Human-specific acceleration of response speed and early spiking over the postnatal period

We used whole-cell patch-clamp recordings to define the postnatal maturation of juvenile human and mouse layer 2/3 (L2/3) neocortical pyramidal neurons through childhood and adolescence (Fig. 1a). We obtained neurosurgically-resected temporal cortex tissue blocks from 24 donors, ranging in age from 9 months to 18 years (Fig. 1b), who were undergoing treatment for removal of an underlying epileptic focus or tumor (see Extended Data Table 1). To delineate human-specific features relative to the far better studied rodent neocortex, we recorded from mice spanning the equivalent developmental window (ages p10-p60) using matched solutions and protocols. We targeted the homologous brain region in mouse, temporal association area^33,36,40,50,51^ (TeA) (Fig. 1a). Based on the strong depth-dependence of physiology in human L2/3 neurons,^27,33,36^ we targeted recordings across the cortical depth of L2/3 in both species (Fig 1a, bottom).

**Fig. 1:**
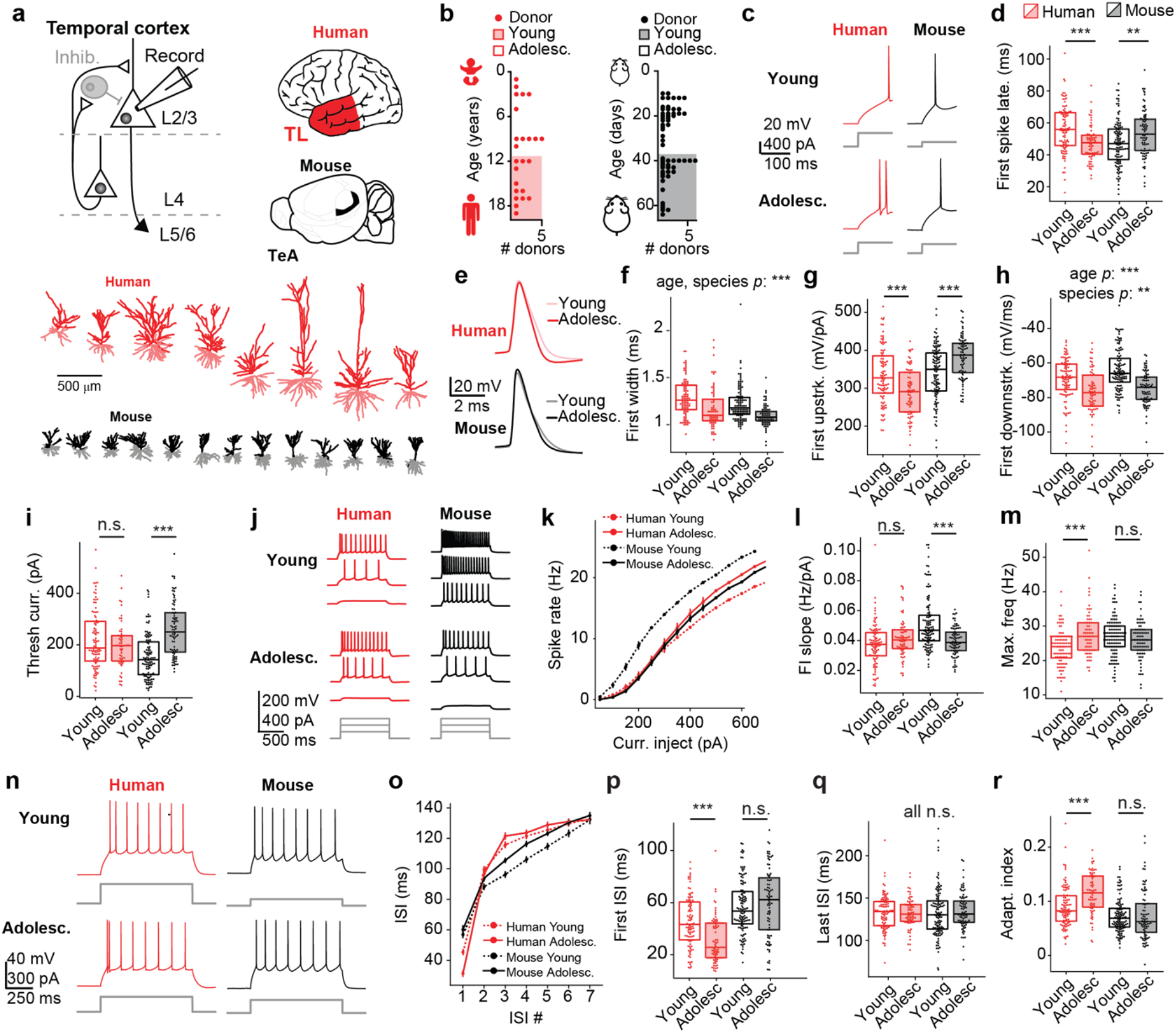
Human-specific changes in intrinsic physiology between early childhood to adolescence favor more rapid responses while preserving input-output relationships. a, Experimental overview (upper): Electrophysiology was measured from neocortical neurons spanning the whole depth of L2/3 in human temporal lobe (TL) from neurosurgical resections and in mouse temporal association area (TeA). Lower: Representative reconstructed morphology of recorded pyramidal neurons across L2/3 in both human and mouse. b, Age distribution of human and mouse donors. c, Representative spike latencies across ages and species. d, Spike latency decreased across human (*p* = 2.28x10^-5^), but increased across mouse (*p* = 0.0034), development. e, Representative spike widths across age and species. f, Spike width decreased across human and mouse development (*p* = 2.58x10^-9^, 0.0002 age, species main effects, two-way ANOVA). g, The action potential upstroke decreased across human (*p* = 2.80x10^-4^), but increased across mouse (*p* = 3.27x10^-5^), development. h, The action potential downstroke decreased across both human and mouse development (*p* = 7.70x10^-13^, 0.0037 age, species main effects, two-way ANOVA). i, The amount of current needed to elicit spiking was stable across human (*p* = 0.50), but increased across mouse (*p* = 1.22x10^-11^), development. j, Representative responses to a range of depolarizing current injections. k, Line plot of the average frequency of spikes elicited in response to different current injections as in (j) across species and ages. l, The slope of the spike-frequency versus current injected relationship in (k), which was stable across human (*p* = 0.052), but decreased across mouse (*p* = 7.78x10^-7^), development. m, Maximum spike frequency across current injections increased across human (*p* = 1.15x10^-4^), but not mouse (*p* = 0.30), development. n, Representative frequency-matched spike trains across species and ages for spike train dynamics analyses. o, ISI length across the spike train. p, The first ISI shortened across human (*p* = 5.47x10^-6^), but not mouse (*p* = 0.27), development. q, The last (i.e., 7^th^) ISI length did not change across development in either species (*p* = 0.62, main effect of age). r, Adaptation index increased across human (*p* = 1.40x10^-5^), but not mouse (*p* = 0.525), development. The number of samples belonging to young human, adolescent human, young mouse, and adolescent mouse, respectively for (d,f,g,h,m,o,p,q,r) are: *n* = 96, 68,115, and 82; for i: *n* = 96, 57, 108, and 81; for (k,l): *n* = 93, 68, 114, and 80. Follow-up pairwise comparisons within a species across ages were performed via two-tailed Mann Whitney U if there were significant species X age interactions in a two-way ANOVA. If there was no significant species X age interaction, significant main effects are reported.

Short spike latencies and fast action potential kinetics are known hallmarks of adult human neocortical neurons^32,37^ and can correlate with intelligence.^35^ We therefore tested if this trait may be an emergent property across critical periods of human development, during which cognition becomes increasingly sophisticated and processing speed increases.^52,53^ Indeed, we found that the onset of spiking (spike latency) accelerated across early childhood to adolescence in human neocortical pyramidal neurons, whereas it decelerated in mouse neurons^8^ (Fig 1c,d). Spike widths became narrower in both human and mouse neurons across development^8,28^ (Fig. 1e,f). However, while the narrowing of spike widths was driven by both a faster upstroke and downstroke in mouse, in humans it was driven primarily by an accelerated action potential downstroke (Fig. 1g,h). Thus, we identify dissociable postnatal regulation of action potential onset timing and kinetics across species.

Shorter spike latencies could be an indicator of upregulated overall excitability. However, we found that excitability was remarkably stable in human neurons across postnatal development. Specifically, a similar amount of current was needed to elicit spiking in both young and adolescent neurons (Fig. 1i) and a similar number of spikes were elicited by a range of suprathreshold current injections (Fig. 1j,k), as quantified by the slope of the frequency-input (FI) curve (Fig. 1l). This was surprising given the changes in spike latency, and yet this result aligns with recent findings in pediatric human L2/3.^27,28^ While the input-output relationship remained the same in human neurons, we found signatures of an expanded dynamic range, as evidenced by an increased maximum firing frequency (Fig. 1m). This suggests an optimization for high-frequency firing to track the rapid synaptic kinetics of adult human L2/3 relative to mouse.^35,37^

By contrast, adolescent mouse neurons required more current to initiate spiking (Fig. 1i) and spiked less when injected with current relative to young mouse neurons (Fig. 1j,k), resulting in a shallower slope of the FI curve (Fig. 1l), with no change in maximal firing frequency (Fig. 1m).^8,10,11^ In line with this age-dependent reduction in excitability, adolescent mouse neurons displayed decreased input resistance and hyperpolarized resting potential relative to young mouse neurons, whereas these properties were stable across human postnatal development (Extended Data Fig. 1).

The primary subthreshold physiological change in adolescent human neurons was pronounced upregulation of the HCN channel-mediated voltage sag. Pharmacological blockade of HCN channels with ZD7288 across ages and species confirmed that this physiological property showed more pronounced postnatal changes in human (Extended Data Fig. 1). These results reveal that the specialized role for HCN channels uncovered in the adult human L2/3 circuit is established during postnatal development.^36^ As sag current can accelerate the kinetics of synaptic responses,^36^ the human-specific upregulation of this current provides further evidence that human-specific optimizations for speed of transmission^35,37^ are developmentally established.

In line with the reduced time to onset of spiking, we found that the first inter-spike interval (ISI) was also markedly reduced in adolescent human neurons when comparing frequency-matched spike trains (Fig. 1n-p). However, the last ISI was comparable in young versus adolescent human neurons (Fig. 1q). We did not observe postnatal changes in first or last ISI in mouse neurons for frequency-matched spike trains (Fig. 1n-q).

This human-specific acceleration of early spiking across development was conserved at higher spike rates. However, at higher spike rates the acceleration extended beyond the first ISI, suggesting that this acceleration of early spiking may be driven by an intracellular signaling process with an intrinsic time-course, rather than a strictly spike-dependent mechanism (Extended Data Fig. 2).

We quantified this change in spike train dynamics by computing a rolling average in the change in ISI length across the train (“adaptation index”; see Methods). Consistent with the selective acceleration of early ISIs, human neurons showed a striking, age-dependent increase in spike rate adaptation (Fig. 1r).^28^ In addition, the baseline adaptation index was higher in human than in mouse neurons (Fig. 1r). Thus, our results reveal that the upregulation of adaptation observed in adult human and non-human primate L2/3 is amplified through postnatal development.^33,42,54^ Moreover, spike rate adaptation has important computational implications, including allowing individual neurons to compensate for changing network activity, statistics or noise levels.^55–58^ Thus, in this case, spike rate adaptation may serve a dual purpose: to promote vigorous neuronal responses without changing the input-output function, and to endow individual neurons with flexible computational properties as the network matures.

We considered if any of our physiological findings interacted with other covariates like cortical depth from pia, neuron morphology, or donor sex. We found that several physiology features showed a striking depth dependence,^27,33,36,54^ with deeper neurons diverging most prominently from mouse (Extended Data Fig. 3). With respect to morphology, we did not find significant correlations between total dendritic path length and key physiology features, with the exception of a significant anti-correlation between input resistance and total dendritic path length in human neurons alone (Extended Data Fig. 4). When we considered sex as a covariate, we found significant sex X age interactions for a subset of physiology features in both species, but trends were broadly conserved across sexes (Extended Data Fig. 5).

## Patch-Seq based linear models identify molecular candidates for human-specific acceleration of early spiking

To generate unbiased insight into the molecular mechanisms underlying the specialized postnatal development of human neurons, we applied a variant of single-cell transcriptomics called Patch-Seq (Fig. 2a; see Methods).^29–31^ We aspirated the contents of each neuron into the recording pipette following physiology recording and sequenced the resulting RNA using next-generation sequencing. This allowed us to establish matched pairs of physiology and gene expression data across species and developmental stages with single-cell resolution. A total of 54 human and 58 mouse neurons spanning developmental stages from early childhood through adolescence passed quality control metrics for both physiology recording and single-cell RNA quantification (see Methods).

**Fig. 2:**
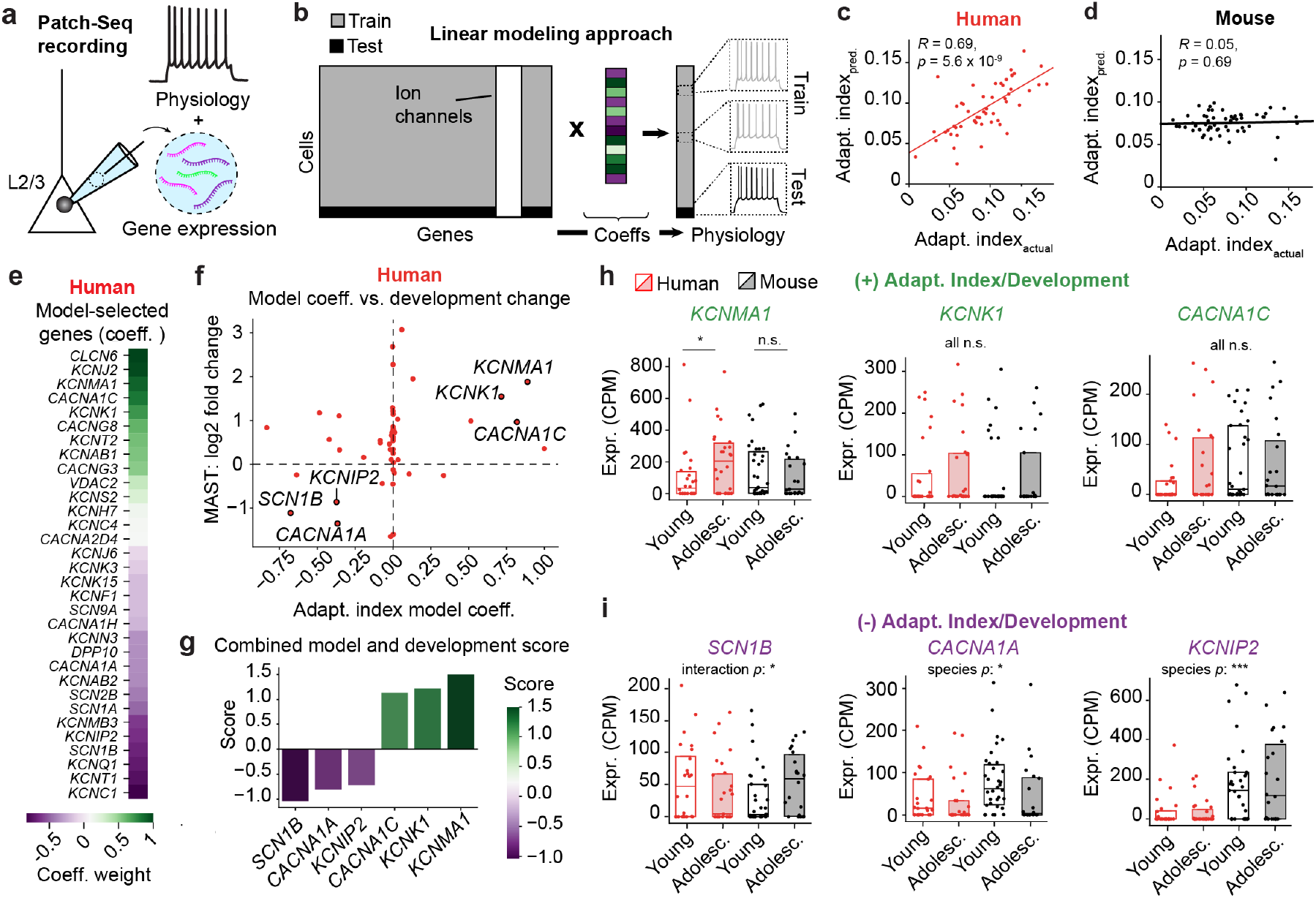
Patch-Seq identifies potential ion channel drivers of developmentally-established, human-specific changes in adaptation. **a**, Schematized Patch-Seq recording, which allows for obtaining paired transcriptomic and physiology data from single neurons. **b**, Linear models were trained to regress physiology (adaptation index) from gene expression (ion channels). Scatter plots denote individual **c,** human (*n* = 54) or **d**, mouse (*n* = 58) Patch-Seq neurons, with the *x*-axis indicating the measured adaptation index and the *y*-axis the predicted adaptation index. Human, but not mouse, model predictions were significantly correlated to measured adaptation index values, quantified via Pearson’s *R*. **e**, Non-zero coefficients assigned to genes by the model, with coefficients scaled to the absolute maximum value across all genes. **f**, Scatterplot displays the assigned adaptation index linear model coefficient (*x*-axis) versus the development fold change (log2fc) quantified with MAST (*y*-axis) for every ion channel gene input into the human adaptation index model. **g**, Sum of development log2fc values and model coefficients scaled to the absolute maximum value. The three most positive and most negative scores are listed, along with the corresponding genes. **h**, Expression of the three most positively-scored genes from (**g**) across development in human and mouse. A two-way ANOVA indicated a trending species X age point interaction for *KCNMA1* (*p*=0.07). Follow-up comparisons indicated that *KCNMA1* expression increased across human (p = 0.028) but not mouse (*p* = 0.50) development. **j**, Expression of the three most negatively-scored genes from (**g**) across development in human and mouse. A two-way ANOVA revealed a significant species X age interaction for *SCN1B* (*p* = 0.045), and main effects of species for both *CACNA1A* (*p* = 0.035) and *KCNIP2* (*p* = 4x10^-6^). Follow-up comparisons did not reveal a significant change in the expression of *SCN1B* across mouse (*p* = 0.09) or human (*p* = 0.21) development. Follow-up comparisons were made via two-tailed Mann-Whitney U. For (**h**,**i**), each dot represents a neuron and the number of neurons belonging to young human, adolescent human, young mouse, and adolescent mouse, respectively, were: *KCNMA1* (*n* = 26, 26, 35, 22), *KCNK1* (*n* = 26, 25, 33, 21), *CACNA1C* (*n* = 25, 25, 33, and 23), *SCN1B* (*n* = 24, 27, 34, 22), *CACNA1A* (*n* = 25, 26, 34, 20), and *KCNIP2* (*n* = 24, 27, 33, 22).

To identify potential drivers of the human-specific acceleration of early spiking that we observe across postnatal development, we fit linear models to human and mouse single-cell Patch-Seq datasets (Fig. 2b). This approach contrasts with most prior work, which abrogated the single-cell resolution in Patch-Seq data by averaging across cell types to detect broader trends^30,31,33,46–48^ (but see^29,59,60^). Although averaging can de-noise single-cell data by limiting artifacts such as gene detection drop-outs, it can also mask biologically meaningful single-cell variation in both physiology and gene expression^61–69^ and consequently preclude identifying mechanisms within a defined cell type.

We separately fit models to human and mouse datasets to determine the extent to which any identified gene-physiology relationships were conserved across species. Because ion channel gene expression is the primary genetic determinant of physiology, we restricted our input gene expression datasets to ion channels (Fig. 2b). This had the added benefit of reducing overfitting that can result from inclusion of a large feature set. We further mitigated overfitting by enforcing model sparsity with L1 regularization.^70^ To fit linear models and quantify model performance, we employed a leave-one-out strategy and computed the correlation between the predicted adaptation index versus the actual measured adaptation index for each held-out test neuron (see Methods).^29^

This approach identified a significant correlation across human Patch-Seq neurons (Fig. 2c), consistent with a predictive model, although we did not see any equivalent correlation in mouse Patch-Seq neurons (Fig. 2d). The same method also fit a predictive linear model of the length of the first ISI to our human Patch-Seq data (Extended Data Fig. 6), but was unsuccessful for spike latency or other single-spike features. The modeling results for first ISI largely mirror those from the adaptation model, leading us to focus primarily on our adaptation findings (but see Extended Data Fig. 6).

We next examined the coefficients assigned to each of the ion channel genes by our model, because the genes that were useful to predict variance in physiology are likely contributors to the underlying cellular mechanisms. Because we used L1 regularization, we would expect predictive (i.e., physiologically-relevant) genes to be assigned non-zero coefficients (i.e., “selected”), whereas relatively unpredictive genes would be assigned coefficients of exactly zero.^70^ Selected ion channels spanned several families, including potassium, sodium, and calcium channel subunits (Fig. 2e; Extended Data Table 2), indicating that an interplay between multiple conductances likely drives postnatal physiological changes.

Ion channels that drive developmental changes in adaptation are expected to be developmentally regulated in addition to predicting variance in adaptation. We therefore plotted model-assigned coefficients against log2 fold-change across development, quantified with the Model-Based Analysis of Single-Cell Transcriptomics (MAST) package in R^71^ (Fig. 2f, see Methods). Genes that occupied regions outside of the *x*-and *y*-axes of this plot show both a relationship to variance in adaptation and changes in expression across development (Fig. 2f).

We combined these two measures to establish a single metric reflecting the relationship of each model-selected gene to adaptation and/or human developmental stage. We scaled both log2 fold-change values and model coefficients for each ion channel gene to the respective absolute maximum value for that measure and then summed these two values. We then selected the three genes with the most positive scores and the three genes with the most negative scores for deeper investigation (Fig. 2g). Each of these genes fell in either the upper right quadrant or the lower left quadrant of our graph, confirming internally consistent relationships to adaptation and human development (Fig. 2f).

Among the three most positively-scored genes, *KCNMA1* showed a significant increase in expression across human, but not mouse, development (Fig. 2h). This hit was surprising because *KCNMA1* encodes the BK-type calcium-activated potassium channel, which is not canonically associated with spike rate adaptation in most neuronal cell types. However, in rat hippocampal pyramidal neurons, BK channels can accelerate spiking early in the train in a highly specific mechanism that is engaged for only 50-100 ms at the start of a spike train. Rapid disengagement of this excitatory mechanism leads to early adaptation of the firing rate.^49^ Our data are consistent with a similar mechanism emerging during adolescent development in human but not mouse L2/3 pyramidal neurons (Fig. 1n-r). Indeed, our linear model of first ISI length assigned one of the most negative model coefficients to *KCNMA1* (Extended Data Fig. 6).

The calcium channel gene *CACNA1C* parallels *KCNMA1* in model coefficients for adaptation and first ISI (Fig. 2e-h; Extended Data Fig. 6), as well as in developmental changes in expression, although developmental changes in *CACNA1C* expression did not reach statistical significance in our Patch-Seq dataset. *CACNA1C* encodes the primary subunit of the L-type calcium channel, which functionally couples with BK channels on the order of single spikes.^72–74^ Our results therefore suggest that human-specific upregulation of BK channels, possibly in synergy with L-type channels, accelerates early spiking in adolescent human neurons.

Among the three most negatively scored genes, we found a species X age interaction in the expression of *SCN1B* (Fig. 2i). Post-hoc comparisons of *SCN1B* expression revealed a trending upregulation across mouse development and a non-significant decrease across human development, suggesting that these bidirectional changes in expression may drive the species X age interaction. The combination of a strong negative weighting of *SCN1B* in our adaptation model (Fig. 2e,f) and a mean decrease in *SCN1B* expression across human development (Fig 3i) suggests that *SCN1B* acts as a negative regulator of adaptation in young human neurons. *SCN1B* encodes the β subunit of Nav1.1 channels and is an important positive regulator of Nav1.1 channel density.^75,76^ Thus, a developmental reduction of *SCN1B* could reduce the overall number of Nav1.1 channels in adolescent human neurons,^76^ leading to sodium channel exhaustion and contributing to spike rate adaptation.^49^

**Fig. 3:**
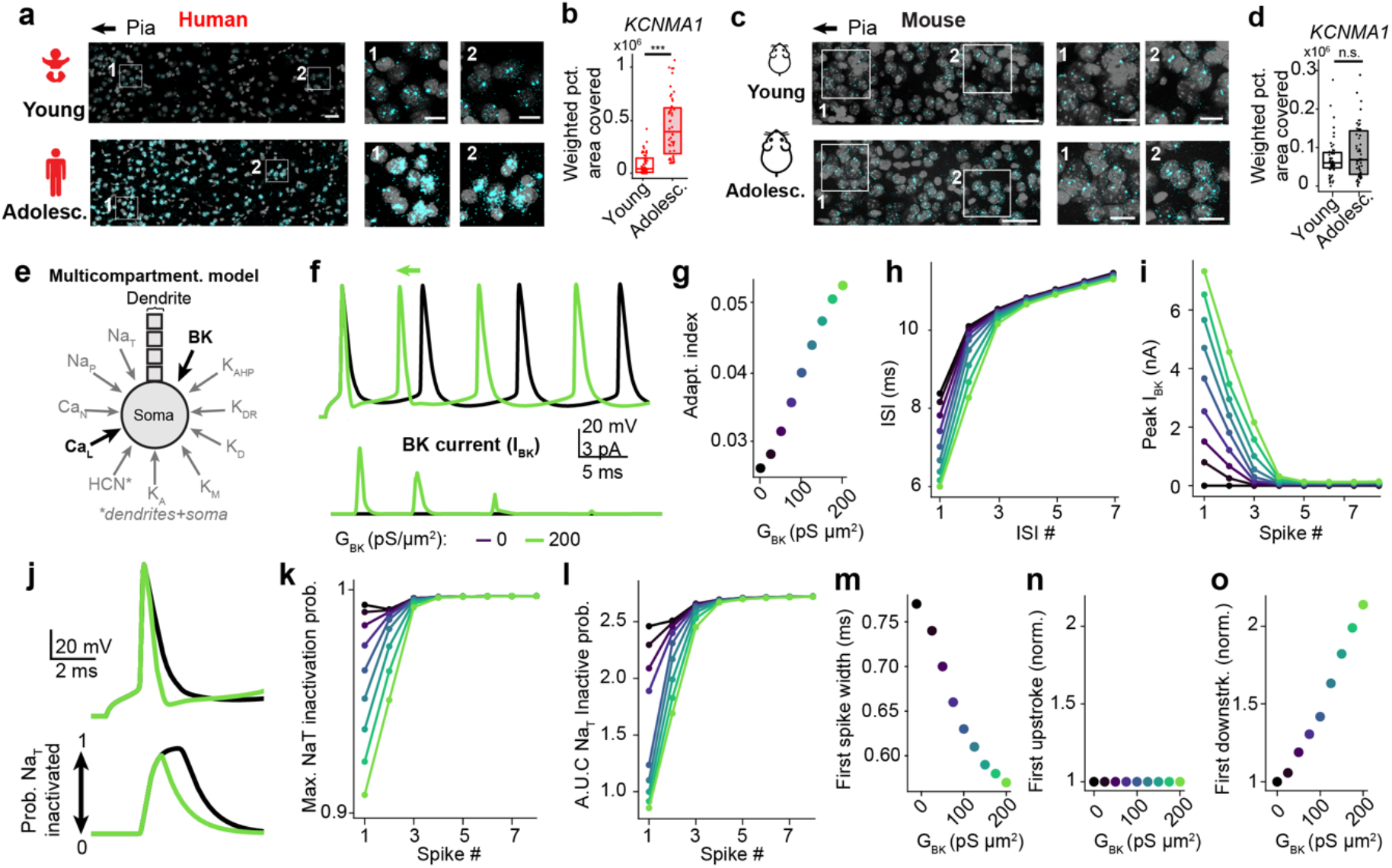
Human-specific postnatal upregulation of BK channels *in situ* and mechanistic involvement in accelerating early spiking *in silico*. a, Representative confocal image of *KCNMA1* (encoding BK channels) RNA labeled *in situ* in young (top) versus adolescent (bottom) human L2/3. **b**, *KCNMA1* expression significantly increased from early childhood to adolescence in humans (*p* = 5.77x10^-7^*, n* = 50 neurons/age, *N* = 4 young human donors, *N* = 3 adolescent human donors). c, Representative confocal image of *KCNMA1* RNA labeled *in situ* in young (top) versus adolescent (bottom) mouse L2/3. **d**, *KCNMA1* expression was stable from early childhood to adolescence in mice (*p* = 0.36; *n* = 50 neurons/age, *N* = 2 young mouse donors, *N* = 2 adolescent mouse donors). **e**, Schematic of multicompartment model and included conductances, with bold indicating conductances corresponding to genes of interest from Patch-Seq based model. **f**, Spike trains from multicompartmental model (top) and corresponding BK current (IBK, bottom). **g**, Adaptation index across different BK conductances. **h**, ISI length across different ISIs and BK conductance levels. **i**, Peak BK current (IBK) across spikes in the train and BK conductance levels. **j**, First spike from multicompartmental model (top) and corresponding NaT inactivation probabilities (bottom). **k**, The maximum NaT inactivation probability attained during each spike in the train. **l**, Area under the curve of the NaT inactivation probability curve for each spike in the train. **m**, Spike width across different BK conductances. As in (**m**) but for upstroke (**n**) and downstroke (**o**). Values in (**o**,**n**) were normalized to the value with no BK current to compensate for differences in magnitude between upstroke versus downstroke to facilitate comparison across these properties.

To complement our targeted physiology-driven approach to transcriptomic analysis, we broadly probed for genes that were differentially expressed across species with MAST (Extended Data Fig. 7). This revealed that human neurons expressed the gene encoding the β4 subunit of the BK channel, *KCNMB4*, at lower levels than mouse neurons. Exclusion of this subunit from BK channels confers fast inactivation kinetics and thus pro-excitatory properties to BK channels in a manner similar to the cross-species difference we observe here.^77^ The observed human-specific, developmentally-regulated form of adaptation may therefore derive in part from lower expression of β4-containing BK channels.

## A developmentally-regulated, BK-mediated mechanism accelerates early spiking

To independently corroborate the human-specific developmental upregulation of *KCNMA1*, we applied multiplexed *in situ* hybridization via RNAscope in another set of samples (see Methods). We identified excitatory neurons across the L2/3 depth in both human and mouse based on expression of an excitatory marker, *SLC17A7,* and quantified *KCNMA1* expression as the optical area of the cell occupied by *KCNMA1* fluorescence, scaled to pixel intensity as quantified by QuPath (“weighted percent area covered”; see Methods). To avoid potential confounds due to species differences in probe efficiency, we examined gene expression within species across development. To ensure even sampling across donors, we pseudo-randomly down-sampled our data to 50 cells per condition, evenly subsampling each donor (see Methods).

In line with our Patch-Seq findings, *KCNMA1* expression increased markedly across human (Fig. 3a,b), but not mouse (Fig. 3c,d), development. Positive and negative controls confirmed that developmental changes were not driven by differences in overall RNA quality, or in the accumulation of the auto fluorescent pigment lipofuscin, which can increase with age in humans^78^ (Extended Data Fig. 8).

To test for underlying mechanisms by which the channel encoded by *KCNMA1*, the BK-type potassium channel, might contribute to spike rate adaptation, we generated a multicompartmental computational model of single neuron physiology (Fig. 3e; see Methods).^49^ Increasing the BK-type potassium current elevated spike rate adaptation in this model (Fig. 3f,g). Consistent with our human physiology recordings, the *in silico* increase in adaptation was driven by a selective shortening of the first ISIs in the train, with later ISIs virtually unaffected (Fig. 3h). In agreement with this, the BK current was greatest in the first spike and then gradually attenuated to near-zero in subsequent spikes (Fig. 3f,i).

We hypothesized that interactions between BK channels and the spike generating mechanism might underlie acceleration of spiking in the presence of BK currents. We found that the level of transient sodium current (Na_T_) was largely agnostic to BK current levels (Extended Data Fig. 9), but the dynamics of Na_T_ channel transitions changed in the presence of BK current (Fig. 3j). Specifically, fewer Na_T_ channels entered the inactivated state and these channels transitioned out of the inactivated state earlier (Fig. 3j). This difference in dynamics was evidenced by a reduction in the maximum inactivation probability (Fig. 3k) and a reduced area under the curve (A.U.C.) of the inactivation probability curve (Fig. 3l) for early spikes in the train. The functional result was that more sodium channels were available earlier to initiate the next spike. Moreover, the modulation of Na_T_ inactivation dynamics was largest for earlier spikes with higher BK current, but in later spikes Na_T_ inactivation dynamics resembled that of models without BK current (Fig. 3k,l).

Completion of the action potential coincided with a switch in sodium channel states from increasing to decreasing occupancy of the inactivated state (Fig. 3j). We therefore hypothesized that BK-mediated acceleration of action potential kinetics might be the key component of this mechanism. Indeed, we found that the presence of BK channels narrowed spike widths (Fig. 3m). Whereas the first action potential upstroke was largely unaffected by the presence of BK channels (Fig. 3n), we found a significant acceleration of the action potential downstroke that scaled with BK conductance level (Fig. 3o). This feature of the model closely paralleled our developmental observations that adolescent human neurons have narrower spike widths due to a selective acceleration of the downstroke (Fig. 1f-h).

## BK channel blockade recapitulates differences in physiology between young versus adolescent human neurons

To causally test this physiological mechanism in human neurons, we bath-applied the BK channel antagonist, Paxilline, during whole cell brain slice recordings. Paxilline reduced spike rate adaptation with a magnitude similar to the changes observed across human postnatal development (Fig. 4a-c and Fig. 1r). This effect was driven by a significant lengthening of the first ISI relative to DMSO controls (Fig. 4d,e), with no change to the last ISI (Fig. 4f). Moreover, there was a strong negative correlation between the change in ISI length and adaptation due to BK channel blockade on a cell-by-cell basis, whereas no relationship was found between the last ISI and adaptation. This result paralleled the relationship between each of these features in our developmental dataset (Extended Data Fig. 10).

**Fig. 4:**
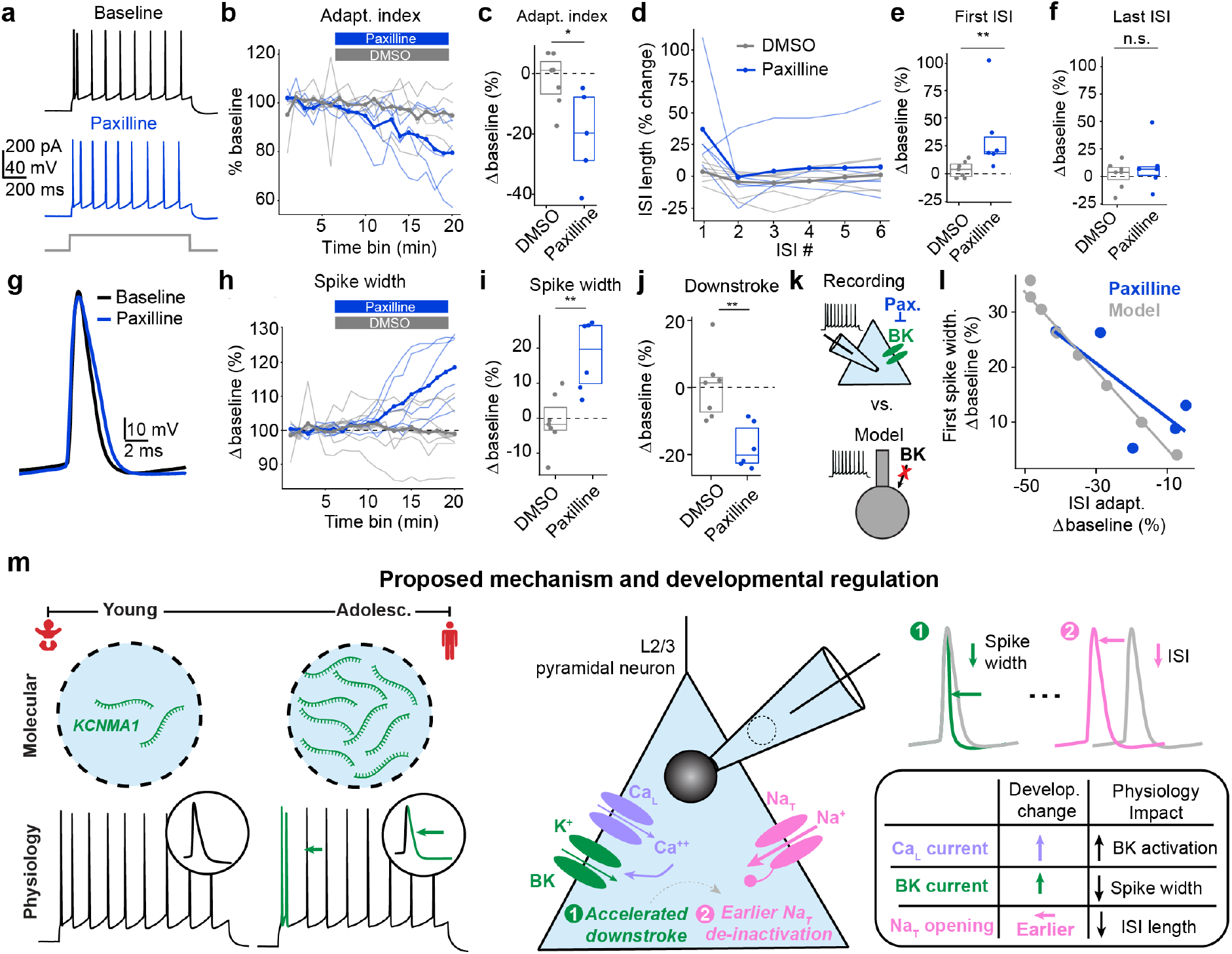
BK channel blockade in human neurons causally validates and extends Patch-Seq predictions. **a**, Representative spike train before (upper) and after (lower) BK channel blockade with Paxilline. **b**, Time course of adaptation index during Paxilline versus DMSO control bath application. **c**, Bath application of Paxilline resulted in a percentage-wise decrease in adaptation index above and beyond DMSO controls (*p* = 0.030; *n* = 7 DMSO, *n* = 5 Paxilline). **d**, Percentage-wise change in ISI length across the spike train after Paxilline versus DMSO control bath application. The first (*p* = 0.008) (**e**) but not last (*p* = 0.731) (**f**) ISI was significantly lengthened after Paxilline application relative to DMSO controls. **g**, Representative first spike before versus after bath application of Paxilline. **h**, Time course of the first spike width during Paxilline or DMSO bath application. **i,** Paxilline bath application significantly lengthened the first spike width (*p* = 0.008) and **j**, slowed the action potential downstroke (*p* = 0.002). Sample sizes for (**d,e,f,h,i,j,l**): *n* = 7 DMSO, *n* = 6 Paxilline. Donor sample sizes for (**b**,**c**,**d**,**e**,**f**,**h**,**i**,**j,l**): *N* = 4 DMSO, *N* = 4 Paxilline. **k**, Two ways of analyzing the impact of BK current on spike width: blockade with Paxilline or removal of BK current from multicompartmental model. **l**, Removal of BK current *in vitro* with Paxilline or *in silico* impacted adaptation index (*x*-axis) and first spike width (*y*-axis) to a similar degree. **m**, Schematized experimental results (left): BK (encoded by *KCNMA1*) expression is selectively upregulated across human development, which results in narrowed spike widths, accelerated early spiking, and upregulated adaptation. Proposed mechanism: Greater expression of BK channels – possibly in synergy with L-type channel upregulation – in adolescent human neurons leads to a faster action potential downstroke, which then allows for earlier de-inactivation of sodium channels. The physiological result is a narrower first spike width and a shorter first ISI.

Further, BK channel blockade significantly widened the first spike^77^ (Fig. 4g-i) and markedly slowed the action potential downstroke (Fig. 4j). Consistent with our *in silico* findings, this pharmacological result provides further evidence that BK channels are activated by a single spike,^72,73^ and thus can shorten ISIs at the immediate onset of spiking.^49^ Upon removal of BK currents either in our computational model or with Paxilline *in vitro* (Fig. 4k), we found a strikingly similar relationship between the degree to which spike width and adaptation were modulated (Fig. 4l). These pharmacology results provide causal evidence that the physiological mechanism predicted by our Patch-Seq-based and multicompartmental model is engaged in human neurons.

Taken together, our experimental and computational results identify a developmentally-regulated mechanism in human neocortical neurons in which BK channels modulate spike width, early spike rate, and adaptation. These features act in concert to facilitate acceleration of early spiking while preserving the input-output function of individual neurons (Fig. 4m).

## Discussion

Here we show that human neurons follow a distinct postnatal physiological trajectory relative to mouse and that this trajectory optimizes human neurons for faster response properties through a developmentally upregulated, BK-channel mediated mechanism. We uncovered this mechanism through a novel linear modeling-based approach to analyzing Patch-Seq data,^29,46,47^ which selected genes based on their ability to explain variability in physiology at single-cell resolution.

Work in humans and rodents links two of our molecular hits, *KCNMA1* (BK channel) and *CACNA1C* (L-type calcium channel), with neurodevelopmental disorders including autism, Fragile X, Timothy syndrome, and developmental delay.^77,79–86^ Moreover, observations of functional coupling and physical clustering of these channels in neurons^72,73^ suggests that our developmentally-upregulated, BK-mediated mechanism likely is enhanced by increased L-type channel expression over the postnatal period. Because the postnatal window that we examine is a critical period during which many neurodevelopmental disorders emerge,^20–24^ disruption of our identified mechanism may contribute to the aberrant neuronal physiology observed in these disorders^20,21^ either through dysfunctional BK or L-type channels. In line with this hypothesis, enhancement of BK channel function has shown therapeutic promise in rescuing Fragile X deficits.^85,87^

A third genetic hit, *SCN1B*, which encodes the β subunit of Nav1.1, showed the inverse relationship to physiology and development as *CACNA1C* and *KCNMA1*. Recent work found that this subunit is subject to regulation by a hominid-specific gene, *LRRC37B*, and that this regulation drives human-mouse differences in neocortical intrinsic excitability.^88^ Our results therefore build on these recent insights to define a new, human-specific role for *SCN1B* in the regulation of postnatal physiological changes.

The mechanism that we uncover closely resembles a BK-mediated mechanism that has been described in rodent hippocampal CA1 pyramidal neurons.^49^ This interchanging of mechanisms across cell types and species is analogous to the recently described upregulation of the HCN-mediated sag current in human relative to mouse L2/3 pyramidal neurons,^36^ as this current is far more prominent in neocortical L5 or CA1 pyramidal neurons in mouse.^89–92^ Taken together, these results suggest a broader evolutionary phenomenon in which the human brain brings existing, modular mechanisms to bear in distinct brain regions and cell types relative to the mouse.^33,35,36^

The disengagement of the BK-mediated mechanism shortly after spiking commences allows human neurons to respond vigorously without changing their input-output properties and leads to a pronounced adaptation of the firing rate. Spike rate adaptation is linked with several distinct cellular and circuit functions, including separating transient signals from noisy background activity,^93^ rescaling gain in response to fluctuating network activity,^55,58^ and performing coincidence detection.^57,94,95^ Each of these functions have also been ascribed to synaptic physiology mechanisms like synaptic depression, feedback, and recurrence.^96^ However, the human L2/3 cortical circuit is characterized by more feedforward and less recurrent synaptic connectivity than mouse L2/3.^39^ We therefore propose that the ability to tune individual neuron response properties through an intrinsic mechanism like spike rate adaptation, instead of through synaptic input, is better-suited for the distinct network architecture of human L2/3 cortical circuits.

Coupling spike rate adaptation with feedforward network connectivity may serve the faster processing speeds^35,37^ and computational abilities of the human brain. Feedforward networks can be faster at transmitting information than recurrent networks,^97^ and spike rate adaptation may be privileged for this kind of transmission. Indeed, adult human L2/3 pyramidal neurons that exhibit spike rate adaptation preferentially connect with pyramidal neurons with low input resistance,^45^ which would be expected to have rapid postsynaptic potential kinetics. This points towards spike rate adaptation as a key mechanism that is engaged at fast nodes in the circuit. Finally, spike rate adaptation can improve performance on a working memory task above and beyond synaptic mechanisms like depression and facilitation in spiking neural networks.^98^ Taken together, these results suggests that spike rate adaptation may be an intrinsic mechanism that acts in concert with rapid synaptic kinetics and feedforward network architecture for fast transmission of information^37^ and improved computational accuracy.^98^

Beyond these immediate biological insights, our findings serve as a critical proof-of-principle demonstration that single-cell ion channel gene expression profiles measured with Patch-Seq contain sufficient signal to uncover biologically-relevant mechanisms within a defined cell type. This opens the door for unbiased identification of candidate, cell-type-specific molecular mechanisms that underlie variability in physiology across other brain regions, cell types, physiological features, developmental stages, and disease states.

One unavoidable limitation of this study is that most of our human neocortical specimens were neurosurgically resected to gain access to an underlying epileptic focus, raising the possibility that disease state and treatment history may influence these results. This concern is mitigated by four primary factors. First, resections are performed to treat focal epilepsy, and the resected tissue lies above the epileptic focus and is not considered part of the primary epileptic zone. Second, we excluded cases where we observed disorganized cortical lamination or synchronous epileptic synaptic events. Third, the changes in spike rate adaptation that we observe across early childhood to adolescence are replicated (although with shallower sampling of juvenile temporal cortex) by a recent study where the majority of tissue was obtained to treat hydrocephalus or tumor.^28^ Fourth, we focus on within-species comparisons to identify changes that vary across human samples, whereas disease pathology is more likely to correlate with cross-species differences.

Predictive models were unsuccessful in fitting other developmentally important physiological changes in human L2/3 neurons, including sag ratio, action potential kinetics, and spike latency in our dataset. Possible reasons for this include that these physiology features could be governed: 1) by genes with non-linearities in their transcription to protein product, 2) by genes that are expressed at low levels or have short transcript lengths and thus may be underrepresented in transcriptomic data,^99^ or 3) by morphology.^40,100^ Our sample size is limited by availability of primary patient samples and the labor-intensive Patch-Seq approach, but future consortium-based approaches building on the success of our single institution study could expand sample sizes. Finally, we uncover a novel BK channel-mediated mechanism, but the experimental conditions optimized for capturing intrinsic physiology are not well suited to isolate the BK channel current itself.

The growing field of human cellular neuroscience is enabling the discovery of distinctive molecular and physiological properties of the human brain. The majority of studies thus far have focused on the adult human brain and recent studies focused on the physiological specializations of the juvenile and the adolescent human brain^27,28^ lack interrogation of underlying mechanisms. Here, we use Patch-Seq, computational modeling, and pharmacology to identify and causally validate a BK-mediated mechanism that drives human-specific physiological changes across a critical postnatal period from early childhood to adolescence. The human-specific developmental changes that we have found in neuronal physiology collectively suggest a postnatal optimization for response speed that is ideally positioned to serve the unique network properties and cognitive abilities performed by the human brain.

## Acknowledgements

This work is supported by funding from a Brain and Behavior Research Foundation Young Investigator Award (to SFO), the Stanford Maternal and Child Health Research Institute (to SFO), the Shurl and Kay Curci Foundation (to SFO), the Foundation for OCD Research (FFOR, to SFO), the Neidig Fund (to SFO), the Tusher Family Stanford Interdisciplinary Graduate Fellowship (to RMK), and the Stanford Summer Research-Amgen Scholars Program (to SSD).

We gratefully acknowledge the Stanford Microscopy Service (confocal imaging) and Stanford Genomics (sequencing of Patch-Seq samples). We thank A. Kreitzer and R.W. Tsien for helpful comments on the manuscript.

## Methods

### Human tissue acquisition and slice preparation

Specimens were obtained from neurosurgical procedures at either Lucile Packard Children’s Hospital or Stanford Hospital. All procedures were performed in strict compliance with protocols approved by the Institute Review Board at Stanford School of Medicine before commencing the study, and all patients provided informed consent.

Following resection, tissue was placed in a sterile, prechilled bottle filled with carbogenated (95% O_2_ / 5% CO_2_) slicing artificial cerebrospinal fluid (ACSF) containing: Sucrose (68 mM), NaCl (55 mM), KCl (2.5 mM), NaHCO_3_ (25 mM), NaH_2_PO_4_ (1.25 mM), HEPES (20 mM), D-Glucose (25 mM), MgSO_4_ (10 mM), CaCl_2_ (0.5 mM), Na-Pyruvate (3 mM), and Na-L-Ascorbate (5 mM). Surgical specimens were quickly transported (within 10-30 minutes) on ice from the operating room to the research facility and immediately processed for sectioning.

### Animals

All animals used in this study were a mix of male and female C57BL/6 mice aged 10 to 65 days old. Breeders were obtained from Charles River Laboratories (Wilmington, MA) and bred in-house at Stanford University in an AAALAC accredited facility on a 12-hour light/dark timed scheduled in a temperature and humidity controlled room with ad libitum access to food and water.

### Brain slice preparation

Euthanasia of animals occurred under deep anesthesia with isoflurane followed by decapitation. All procedures were performed in strict compliance with a protocol approved by The Stanford University Animal Care and Use Committee. Brains were quickly extracted following decapitation and placed in chilled carbogenated dissection ACSF. Brain slice preparation was similar for both mouse and human specimens.

Specimens were sectioned using a Leica VT1200S Vibratome in chilled carbogenated slicing ACSF at 300 or 350 μM sections for mouse or human, respectively. Mouse brains were sectioned in the coronal plane while human specimens were sectioned through the long axis of the gyrus to obtain all six cortical layers. Brain slices were then placed in a submersion holding chamber for 10 minutes filled with warmed 34°C incubation ACSF^33^ containing: NaCl (100 mM), KCl (2 mM), NaHCO_3_ (25 mM), NaH_2_PO_4_ (1.25 mM), HEPES (20 mM), D-Glucose (25 mM), MgSO_4_ (2 mM), CaCl_2_ (2 mM), Na-Pyruvate (3 mM), and Na-L-Ascorbate (5 mM). Slices were transferred to a submersion holding chamber filled with 25°C incubation ACSF until used for electrophysiological recording.

Brain slices were placed in a submersion recording chamber continuously perfused at 2-3 ml/min with warmed 30°C recording ACSF containing: NaCl (124 mM), KCl (2.5 mM), NaHCO_3_ (24 mM), NaH_2_PO_4_ (1.2 mM), HEPES (5 mM), D-Glucose (12.5 mM), MgSO_4_ (2 mM), and CaCl_2_ (2 mM). Slices were visualized with a Scientifica SliceScope Pro 3000 microscope with infrared differential interference contrast (IR-DIC) optics and a 40x water immersion objective. Patch pipettes (3-4 MΩ) were filled with internal recording solution containing: K-gluconate (110 mM), KCl (12 mM), HEPES (10 mM), EGTA (0.2 mM), ATP-Mg (1 mM), GTP-Na_2_ (0.3 mM), Na_2_-Phosphocreatine (10 mM), Glycogen (20 μg/mL), RNase inhibitor (0.25 U/μL), Biocytin (13.4 mM), and Alexa 488 (0.05 mM). The pH was adjusted to 7.25 with KOH.

### Physiology recording and quality-control

Whole-cell recordings were acquired using a Multiclamp 700B amplifier with custom data acquisition software MIES (https://github.com/AllenInstitute/MIES/) written in Igor Pro. Electrical signals were digitized at 50 kHz with a National Instruments PCIe-6343 digitizer and lowpass filtered at 10 kHz. Upon attaining a whole-cell current clamp configuration, the signal was bridge balanced and pipette capacitance was compensated. The liquid junction potential was calculated to be -14 mV using LJPcalc and was not corrected. Pyramidal cells across the cortical depth were targeted based on morphology under DIC. Pyramidal cell morphology was again confirmed after recording based on Alexa fills with a 540/605 nm excitation/emission filter set. Alexa fills were used to measure distance from pia after recording. Human neurons ≥ 1500 µm and mouse neurons ≥ 400 µm from pia were excluded post-hoc.

All sweeps from individual recordings were visually inspected and sweeps with noise or synaptic events were excluded. All recordings had access resistance <20 MΩ. All neurons in the dataset had V_rest_ > -55 mV. All human neurons had R_in_ < 300 MΩ consistent with prior work in pediatric human samples.^27,28^ All mouse neurons had R_in_ < 500 consistent with prior work in the juvenile period in mice.^8^

Spike detection was performed with custom Python scripts based on the Allen Institute IPFX package. Briefly, spikes were required to cross 0 mV with at least 20 mV/ms slope, and to revert to 0 dV/dt between spikes. A maximum of 5 ms was allowed between the spike threshold and peak. All spikes in the dataset had a peak > 25 mV.

### Physiology protocols

We used three current stimulus protocols to characterize the intrinsic physiology of human and mouse neurons:

1. A series of 1 s current steps from 50 pA to 1000 pA in +50 pA increments
2. A linearly increasing current across a 16 s window, with the minimum current 0 pA and the maximum current per step ranging from 50 pA to 210 pA with +20 pA steps (Ramp)
3. A series of 1 s current steps from -150 pA to 50 pA in +20 pA increments

### Physiology features

We computed spike latency, single-spike properties (i.e., spike width, upstroke, and downstroke), FI slope, maximum frequency, adaptation index, and ISI length from the first protocol. Spike latency was defined as the time between the start of current injection and the peak of the first spike. Spike width was defined as the time between half-height on the upstroke and the half-height on the downstroke, where height is defined as the difference in mV from the action potential trough to peak. The action potential upstroke was defined as the maximum dV/dt value in mV/ms between the action potential threshold and the action potential peak, and the downstroke as the minimum dV/dt value between the action potential peak and the trough. The adaptation index was computed as the average change in ISI length across the train, normalized to ISI length, according to the following formula:

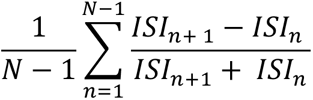

To calculate the minimum amount of current needed to elicit a spike, we extracted the current injected at the time of the first spike threshold from the ramp protocol (2).

We computed the following subthreshold properties from the third protocol: resting potential, input resistance, and sag ratio. Traces with action potentials either during the baseline, current injection, or inter-sweep interval were excluded. Resting potential was calculated as the average of the baseline before current injection across sweeps. Input resistance was calculated as the slope of the relationship between injected current and the steady state voltage deflection across all current injections that passed quality control. Sag ratio was computed as the fraction of the steady-state versus peak input resistance.^101^

### Physiology data analysis and statistics

All data analysis was performed with custom scripts in Python. For each physiology feature, we ran a two-way ANOVA using the statsmodels.formula.api.ols function. We followed up on significant species x age interactions with two-tailed Mann-Whitney U tests.

Physiology boxplots, where the middle line is the mean and the box edges are the first and third quartiles, were disaggregated by species and age, depth, or sex and overlaid with a seaborn.stripplot.

To examine co-variation between depth and age, we generated a binned average across depths and ages for a given physiology feature. We applied the gaussian_filter function as part of the scipy.ndimage package with σ = 1 and plotted the resulting values as a heatmap using the seaborn.heatmap function.

### Patch-Seq sample collection and sequencing

Patch seq samples were processed as previously described (Lee et al., 2021). Prior to data collection for these experiments, all surfaces were thoroughly cleaned with RNase Zap, and as needed RNase Away. At the end of Patch-seq recordings, negative pressure (−20 mbar) was applied through the pipette for ∼5 min after which the nucleus was extracted by very slow pipette withdrawal with higher negative pressure (−70 to -100 mbar). The pipette was removed from the recording chamber and the contents of the pipette were expelled into a PCR tube containing lysis buffer (Takara, 634894). Patch-seq sample tubes were held on dry ice in a benchtop plexiglass enclosure throughout the recording session to ensure collected samples remained free of RNase and DNase contamination. Sample tubes were then transferred to -80C for storage until further processing.

Single-nucleus samples were isolated and lysed using the lysis buffer from the SMARTer Ultra Low Input RNA Kit for Sequencing - v4 (Takara Bio). The cDNA was amplified for 18 cycles to ensure complete 5′ end coverage and unbiased transcript representation. Subsequently, 0.150 ng of cDNA samples (>1,000 bp in size) was used for library preparation with the Illumina Nextera XT kit following the standard protocol. Uniquely dual-indexed libraries were pooled and sequenced on a DNBSEQ-T7 platform (MGI/Complete Genomics). All samples, including positive and negative controls, were treated identically.

### Patch-Seq Transcriptomics data alignment and count quantification

We indexed reference genomes for each species using the STAR command genomeGenerate. For human samples, the GRCh38 GenBank human genome file GCA_000001405.15_GRCh38_no_alt_analysis_set.fna was downloaded from https://ftp.ncbi.nlm.nih.gov/genomes/all/GCA/000/001/405/GCA_000001405.15_GRCh38/seqs_for_alignment_pipelines.ucsc_ids/. This genome was indexed using the following annotation file: gencode.v36.annotation.gtf, downloaded at https://www.gencodegenes.org/human/release_36.html. For mouse samples, the GRCm38.p3 GenBank mouse genome file GCA_000001635.5_GRCm38.p3_no_alt_analysis_set.fna was downloaded from https://ftp.ncbi.nlm.nih.gov/genomes/all/GCA/000/001/635/GCA_000001635.5_GRCm38.p3/seqs_for_alignment_pipelines.ucsc_ids/, and indexed with gencode.vM4.annotation.gtf from https://ftp.ebi.ac.uk/pub/databases/gencode/Gencode_mouse/release_M4/.

The following analysis steps were the same for both species. Reads were mapped to the genome and the transcriptome using STAR in paired-end read mode, after adaptor trimming with fastp. Reference transcriptomes were produced from the reference genome and annotation files using gffread. Reference transcriptomes were used to compute an expression counts matrix from the reads mapped to the transcriptome via salmon.

### Patch-Seq linear modeling

For linear modeling experiments, we first filtered the dataset on genes that were expressed at non-zero levels in at least 10% of the cells in the dataset. Raw counts + 1 were normalized to the total RNA expressed on a per-cell basis and then log-transformed.

We fit L1-regularized linear models using sklearn.linear_model.LassoCV, allowing for an intercept (fit_intercept = True). In the outer loop, we used a leave-one-out approach such that we trained models to predict physiology from gene expression data on all Patch-Seq neurons for a given species, and tested on the excluded Patch-Seq neuron. In the inner loop, we used 10-fold cross-validation on the training data to pick the regularization strength (alpha) for L1-regularized, or LASSO,^70^ regression.

We chose LASSO regression because it enforces sparsity of model fits, such that high-magnitude coefficients are penalized and certain coefficients are dropped to zero. This mitigates overfitting concerns and reduces the set of features to a set that are ‘selected’ as useful to the model, improving model interpretability.

To determine the relative contribution of genes in the dataset while retaining the sign (e.g., positive or negative) of their relationship to physiology, we scaled coefficients to the absolute maximal coefficient value. To uncover robust gene-physiology relationships, we filtered our gene set to those that were assigned same-signed, non-zero coefficients on ≥90% of model iterations.

### Differential gene expression

Differential gene expression across development was assessed via the Model-Based Analysis of Single-Cell Transcriptomics (MAST) package in R. This relies on a hurdle model to estimate differential gene expression in zero-inflated single-cell transcriptomic data.^71^ MAST was run on transcript-per-million (TPM) values, which were extracted from the salmon output, per the package recommendations.

We computed a combined score based on the MAST-computed differential expression and the assigned model coefficient, with each measure scaled to the absolute maximum value, such that these two measures occupied a similar range. We selected the genes with the three most negative or the three most positive scores for follow-up statistical testing to reduce the number of statistical comparisons.

For direct statistical comparisons, we filtered out outliers on a per-gene basis according to the standard exclusion criteria, which required that expression values be within 3 standard deviations of the mean.

### Tissue processing for histology

For mouse experiments, mice were injected with a lethal dose of Ketamine-HCl. Upon loss of the toe pinch reflex, mice were transcardially perfused with 10 mL 1x RNAse free PBS, followed by 10 mL 4% paraformaldehyde in RNAse free PBS. Brains were dissected out and immediately post fixed for 24h in 4% paraformaldehyde in RNAse free PBS. Transverse human tissue blocks were prepared from neurosurgical resections as previously described and drop fixed in 4% PFA in RNAse-free PBS for 24h.

Subsequent processing was the same for mouse and human tissue. After 24h fixation in 4% PFA, tissue was dehydrated through a series of increasing sucrose concentrations: 10%, 20%, and 30% sucrose in RNAse-free PBS. Tissue was moved to a greater concentration sucrose solution upon sinking. Tissue was then frozen in Tissue-Tek O.C.T. embedding media on dry ice. OCT-embedded tissue blocks were cut at 20 μm thickness using a Leica CM1950 Cryostat set to -20°C. Mouse sections were cut in the coronal plane and human sections were cut transversely along the gyrus.Sections were mounted onto glass slides and stored at -80°C with desiccant and enclosed in a plastic bag until use.

### Multiplexed fluorescent in situ RNA hybridization (RNAscope)

Fluorescent *in situ* RNA hybridization was performed using the Advanced Cell Diagnostics (ACD) RNAscope Multiplex Fluorescent Reagent Kit v2 (323100). Sample preparation followed the protease-based pretreatment protocol for fixed-frozen tissue in the RNAscope Multiplex Fluorescent Reagent Kit User manual, with the following modifications: After the initial bake step, slides were post-fixed for 90 minutes at room temperature (rather than 15 minutes at 4°), and two additional 30-minute baking steps were added after tissue dehydration and target retrieval. These steps were added after pilot experiments revealed that these measures prevented tissue detachment in pediatric human tissue samples. Mouse and human samples underwent the same modified pretreatment protocol.

All probes were purchased from Advanced Cell Diagnostics (Hayward, CA) and were the following: Hs-*SLC17A7* (#415611-C2), Hs-*KCNMA1* (#852191-C3), Mm-*Kcnma1* (#476251), and Mm-*Slc17a7* (#416631-C3). C2 and C3 probes were diluted in either a pre-diluted C1 probe or using probe diluent (#300041) at 1:50.

Fluorophore dilutions were empirically determined in both mouse and human samples to maximize signal-to-noise, by preventing under-detection or oversaturation of RNA signal. For human, we diluted TSA Vivid 520 (paired with *SLC17A7*) 1:2000 and TSA Vivid 570 (paired with *KCNMA1*) 1:5000. For mouse, we diluted TSA Vivid 520 (paired with *SLC17A7*) 1:5000 and TSA Vivid 520 (paired with *KCNMA1*) 1:15,000. We ran positive and negative control probes to determine if increased *KCNMA1* signal across human development was driven by overall differences in RNA quality (which would be expected to have higher variability than in the controlled setting for mouse experiments) and/or greater accumulation of the auto-fluorescent pigment lipofuscin with age (reported specifically in humans^78^). We used the following positive control probes, provided premixed at 1:50 dilution: *POLR2A* (C1), *PPIB* (C2), and *UBC* (C3). Negative control probes for C1-C3 were targeted against the bacterial gene *dapB*.

### Image acquisition

Images were acquired on a Zeiss LSM980 inverted confocal microscope with a 63x objective. Z stacks were acquired at 0.45 μm thickness. The same imaging parameters were used for each probe across samples and ages within a species. Imaging parameters were matched between gene of interest probes and negative control probes on a per-channel basis.

Tile scans were used to capture the L2/3 cortical depth and stitched together using the Zeiss ZEN microscopy software. Final analyzed images were maximum-projected Z-stacks. The “Tile Blend” function was used to remove occasional tiling artifacts for display of DAPI images, but all quantification was done on raw images. Care was taken to ensure that segmentation was not impacted by mild tiling artifacts like differential illumination at the boundaries.

### Multiplexed fluorescent in situ RNA hybridization (RNAscope) analysis

Analysis of RNAscope images leveraged the QuPath software and custom scripts in Python. DAPI staining was used to segment individual cells via the “Cell detection” function in QuPath. Excitatory cells were classified according to nuclear *SLC17A7* expression using “Create single measurement classifier.” Custom thresholds were set for each sample based on choosing a value in-between bimodal peaks in expression in a log-transformed fluorescence distribution. To identify fluorescent puncta, we used the “Subcellular detection” function. Thresholds for classifying fluorescence puncta were tailored to each probe based on visual inspection of identified detections and then matched across samples within a given species. This function allowed for both identification of the optical area covered by fluorescent detections, in conjunction with their fluorescence intensity. Cell classifications and fluorescent subcellular detections were visually inspected for each sample.

To quantify expression levels, we extracted the optical area and mean intensity of all subcellular detections. We computed a weighted sum of the optical area covered by detections where the optical area for a given detection was scaled to the mean fluorescent intensity. This weighted sum was then normalized to the total cell area (computed by QuPath based on DAPI segmentation) to generate a “Weighted Percent Area Covered” metric. Normalization to cell area allows for correcting for potential differences in cell size across samples and ages.

To facilitate comparison across our Patch-Seq and RNAscope datasets, which varied in number of cells by an order of magnitude, we randomly down-sampled our RNAscope data to 50 cells/group. To ensure even contributions of each donor to the group average, we evenly subsampled each donor. This had the added benefit of mitigating the tendency for *p*-values to dramatically shrink as a function of sample size, precluding practical, biological interpretation.^102^

To test if any observed changes across human development could be due to systematic differences in RNA quality, we normalized our *KCNMA1* gene expression to the expression of positive control genes on a per-donor basis. To test if the accumulation of the autofluorescent pigment lipofuscin with age might explain expression differences, we computed a “lipofuscin-corrected” metric where the average Weighted Percent Area Covered of detections from negative control samples was subtracted from single-cell *KCNMA1* expression on a per-donor basis. We chose to subtract to avoid dividing by zero, as several negative control samples had zero subcellular detections.

### Pharmacology

*ZD7228*. We bath-applied 25 µm ZD7228, diluted in recording solution, in young and adolescent mouse and human neurons. We continuously ran protocol (3) with a 4s inter-sweep interval. We visually inspected all traces and excluded those with unstable membrane potential or synaptic events that would preclude accurate calculation of the sag ratio.

*Paxilline*. We bath-applied 5 µm Paxilline diluted in recording solution^103^ (with the exception of one pilot cell in the dataset, where we applied 10 µm Paxilline). Paxilline was dissolved in DMSO to make a 50 mM or 100 mM stock solution (for one cell), and then diluted 1:10,000 in recording solution to reach the final working concentration of 5 µm or 10 µm, respectively. For control DMSO washes, DMSO was dissolved in recording solution at 1:10,000 to match the DMSO dilution of Paxilline washes.

All neurons in the dataset demonstrated stable ISIs across at least a 5-minute baseline prior to applying drug. All timeseries were subset on the last five minutes of baseline before wash-on of the drug or DMSO. The stimulus protocol consisted of 2-3 1s depolarizing current injections (with the exception of one pilot cell with 500 ms injections). Current injections were chosen to elicit ∼8-12 spikes. Physiology features were extracted and averaged into 1min bins. Adaptation was computed only for sweeps that elicited at least 5 spikes, and adaptation was only computed for cells that had ≥5 spikes in at least 50% of sweeps within each bin. Computed values were normalized to the average value across the baseline period to obtain percentage-wise changes in physiology features after drug or DMSO application.

### NEURON model

We built a multicompartment model in NEURON^104^ based on a model described in Gu et al. 2007.^49^ In brief, the five-compartment model consisted of an isopotential soma and four dendritic compartments. The model included 11 conductances: B_K_, K_AHP_, K_DR_, K_D_, K_M_, K_A_, HCN, Ca_L_, Ca_N_, Na_P_, Na_T_. All conductances were restricted to the soma, except for HCN which was added uniformly to the dendritic compartment. A key component of this model is rapid coupling between BK channels and voltage-gated calcium conductances, which is implemented by a core-shell calcium model. In this model, calcium first enters a shell compartment where it can couple with and activate BK channels. Diffusion of calcium to a second shell allows for the activation of a second, slower calcium-activated potassium conductance, K_AHP_, which has an inhibitory effect on spiking.

Simulations were run in Python with a 0.01 ms time-step and Crank-Nicolson numerical integration for second-order accuracy in time. Prior to delivery of a current pulse to elicit spiking, a bias current was injected to hold the model at -70 mV for 1000 ms with the NEURON function SEClamp. We injected 600 pA for 100 ms as a point process. We applied the same functions used to analyze our physiology data to quantify various physiology features (see *Physiology features*). To compute the adaptation index, we considered the first 8 spikes in the train. To compare the relative impact of BK current on different physiology features with varying magnitudes (e.g., upstroke versus downstroke), we normalized the raw physiology values to the value in the absence of BK current.

### Morphological reconstructions

A horseradish peroxidase (HRP) enzyme reaction using diaminobenzidine (DAB) as the chromogen was used to visualize biocytin-filled neurons after physiology recordings. Slices were mounted and imaged on a Zeiss AxioImager Widefield Fluorescence Microscope, using brightfield and the 10x object. We acquired 1.78 µM Z-stacks to capture the full extent of the neuron and its processes. To obtain a total dendritic path length, neurons with readily apparent processes and a full apical dendrite were traced with a radial step size of two using Simple Neurite Tracer in the Neuronanatomy plug-in of Fiji.

## Extended Data

**Extended Data Fig. 1:**
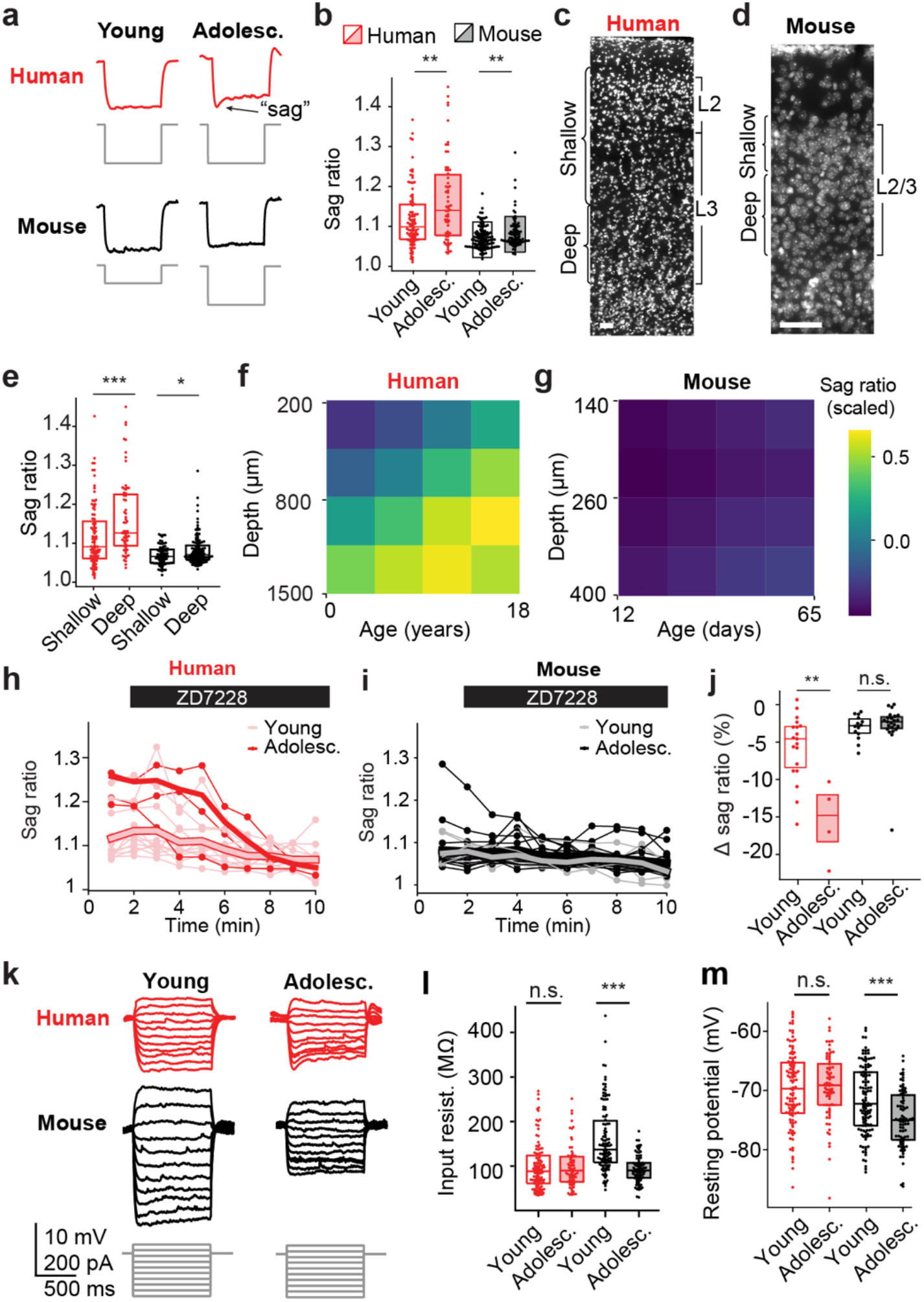
Human neocortical neurons show pronounced upregulation of the HCN-mediated sag current but do not display changes in subthreshold properties canonical to mouse postnatal development. **a**, Example traces showing responses to hyperpolarizing voltage steps across species and ages, with the h-current-mediated voltage sag labeled for one. **b**, Sag ratio increased across both human (*p* = 0.0096) and mouse (*p* = 0.0092) development, though the magnitude of the change is larger in human development. Representative DAPI image of L2/3 with shallow and deep cut-offs denoted for human (**c**) and mouse (**d**). **e**, Sag ratio displayed depth-dependence in human (*p* = 8.40x10^-4^) and to a lesser extent mouse (*p* = 0.026). Sag ratio as a function of depth and age in human (**f**) and mouse (**g**). Sag ratio time course during wash-on of HCN inhibitor, ZD7228 for young and adolescent humans (**h**) or mice (**i**). **j**, The percentage-wise decrease in sag ratio after ZD7228 wash-on increases across human (*p* = 0.0041), but not mouse (*p* = 0.32), development. **k**, Representative traces after subthreshold current injections **l**, input resistance was stable across human (*p* = 0.32), but decreased across mouse (*p* = 3.52x10^-13^), development. **m**, Resting potential was stable across human (*p* = 0.068), but hyperpolarized across mouse (*p* = 1.78x10^-4^), development. Sample sizes for young human, adolescent human, young mouse, and adolescent mouse, respectively (**b**,**l**,**m**): *n* = 96, 60, 114, and 81; for (**h**,**i**): *n* = 8, 4, 9, and 11; for (**j**): *n* = 19, 5, 14, and 23. Sample sizes for shallow human, deep human, shallow mouse, and deep mouse, respectively (**e**-**g**): *n* = 93, 63, 78, and 117. For boxplots, each dot represents a neuron, the center line represents the median, and the box edges represent the 1^st^ and 3^rd^ quartiles.

**Extended Data Fig. 2:**
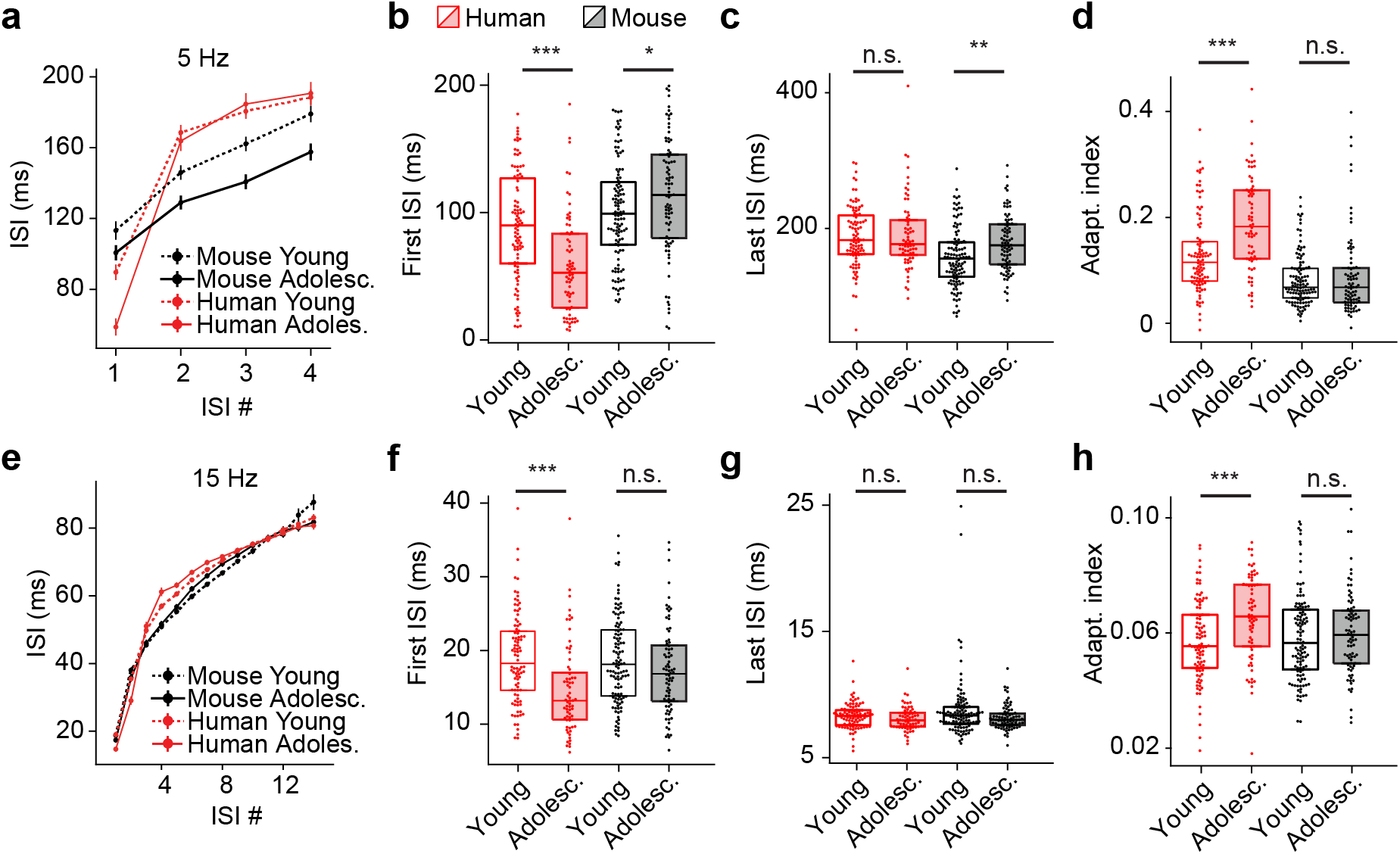
Adaptation across different spike rates. **a**, ISI length for 5 Hz spike trains across species and ages. For 5 Hz spike trains: **b**, first ISI decreased across human (*p* = 3.64x10^-6^), but increased across mouse (*p* = 0.033), development, **c**, last ISI was stable across human (*p* = 0.71), but increased across mouse (*p* = 0.0049), development, **d**, adaptation increased across human (*p* = 9.98x10^-6^), but not mouse (*p* = 0.63), development. **e**, ISI length for 15 Hz spike trains across species and ages. For 15 Hz spike trains: **f**, first ISI decreased across human (*p* = 6.20x10^-8^), but not mouse (*p* = 0.10), development, **g**, last ISI did not change across human (*p* = 0.064) or mouse (*p* = 0.073) development, **h**, adaptation increased across human (*p* = 1x10^-5^) but not mouse (*p* = 0.34) development. The number of samples belonging to young human, adolescent human, young mouse, and adolescent mouse, respectively for (**a**-**d**): *n* = 96, 68, 115, and 82; for (**e**-**h**): *n* = 94, 68, 114, and 80.

**Extended Data Fig. 3:**
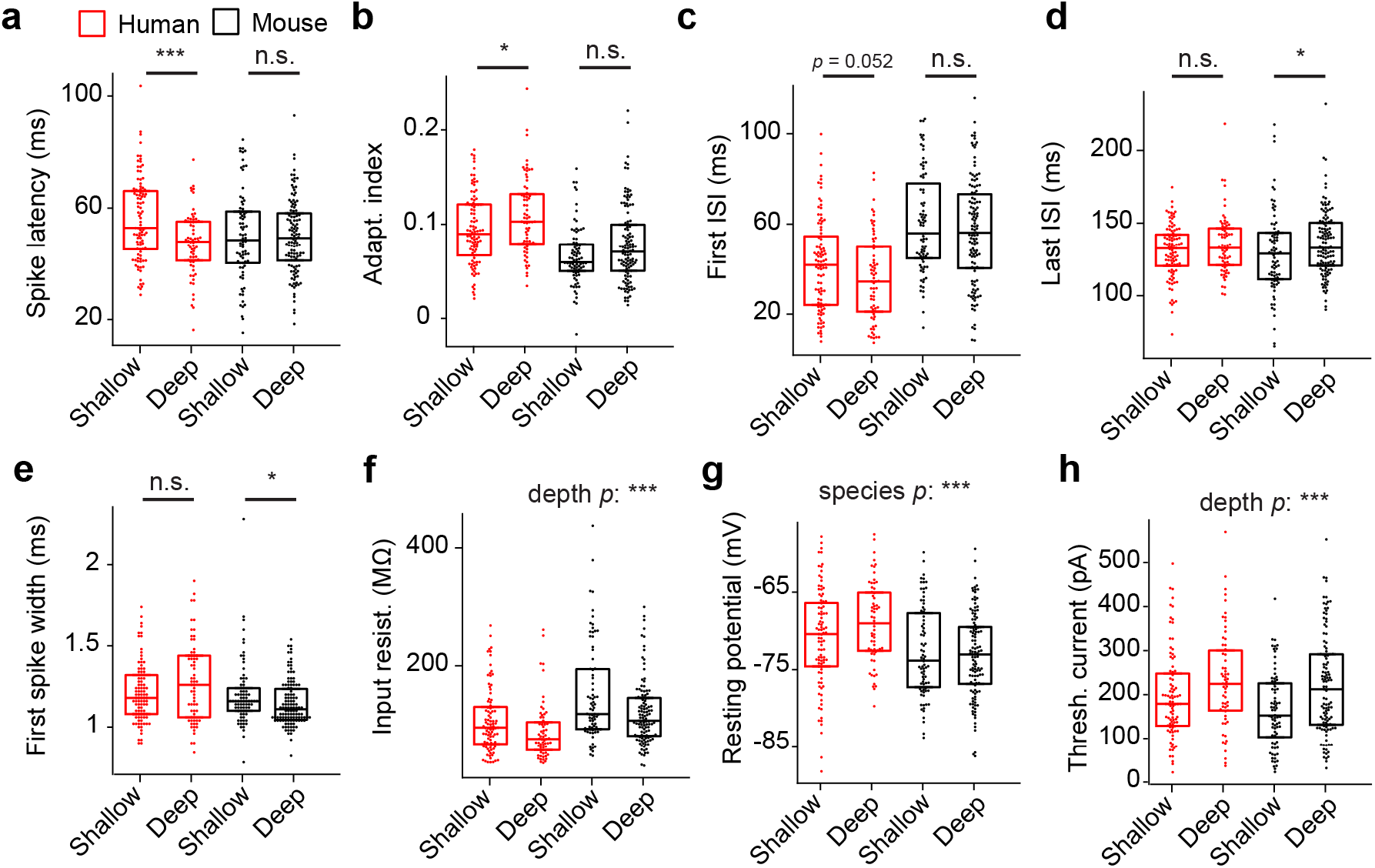
Depth dependence of physiology features. **a**, Spike latency decreased across the depth of human L2/3 (*p* = 0.0006), but did not significantly change across mouse L2/3 (*p* = 0.91). **b**, Adaptation increased across human (*p* = 0.038), but not mouse (*p* = 0.088), L2/3. **c**, First ISI showed a trending decrease across the human L2/3 depth (*p* = 0.052), but no change across mouse L2/3 (*p* = 0.41). **d**, Last ISI was stable across the human L2/3 depth (*p* = 0.34), but increased across mouse L2/3 (*p* = 0.041), **e**, Spike width was similar across human L2/3 (*p* = 0.12), but decreased across mouse L2/3 (*p* = 0.016), **f**, Input resistance decreased across both human and mouse L2/3 (*p* = 0.00010, main effect of depth), **g**, Resting potential was depolarized in human relative to mouse, a difference that was accentuated with depth (species main effect *p* = 4x10^-6^). **h**, Deeper neurons require more current to initiate spiking (main effect of depth, *p* = 4.6x10^-5^). The number of samples belonging to shallow human, deep human, shallow mouse, and deep mouse, respectively (**a**-**e**): *n* = 97, 67, 79, and 118; for (**f**, **g**): *n* = 93, 63, 78, and 117; for (**h**): *n* = 91, 62, 76, and 113. *R* values correspond to Pearson’s coefficient.

**Extended Data Fig. 4:**
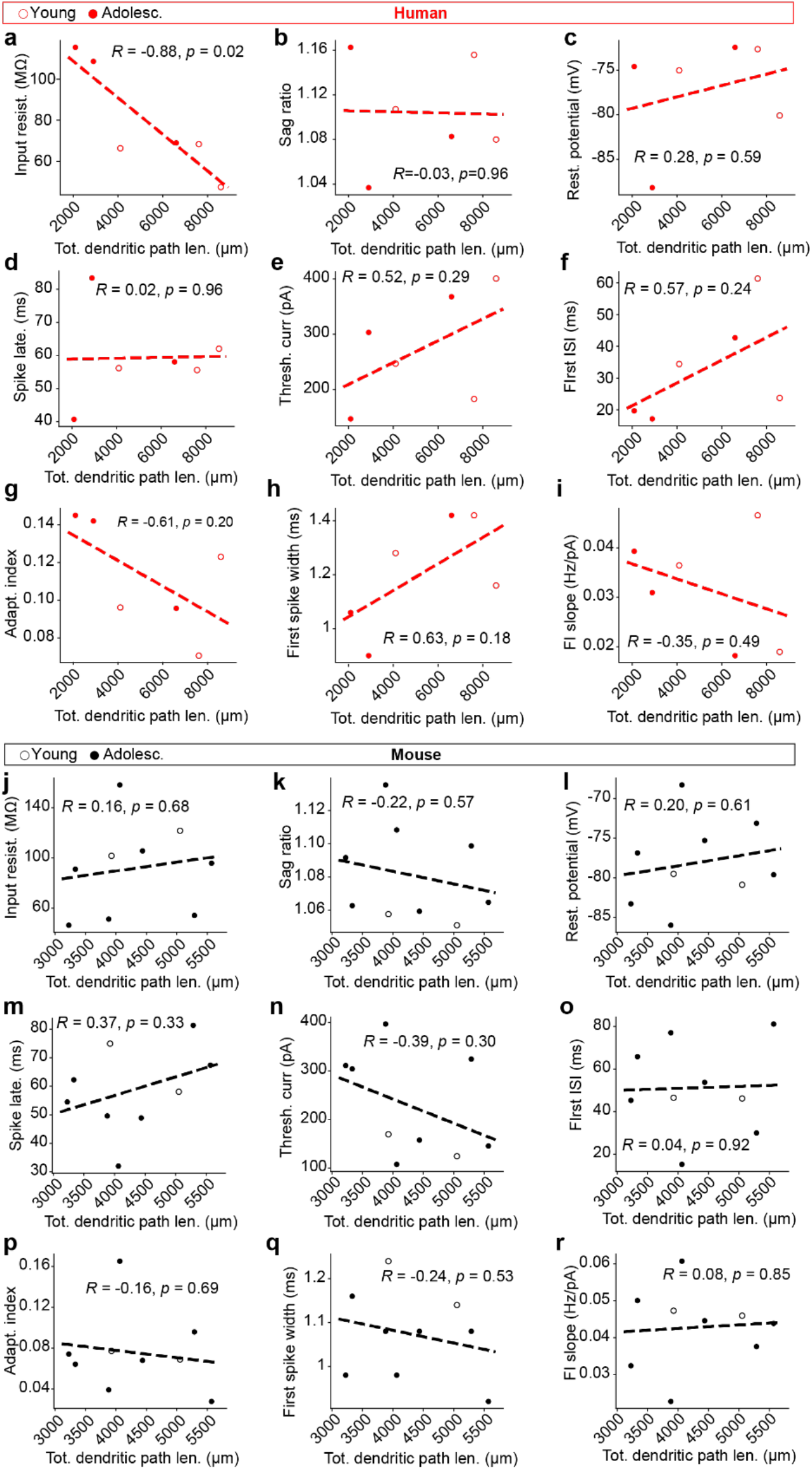
Developmentally-regulated intrinsic physiology properties versus total dendritic path length. Correlation between total dendritic path length and: **a**, input resistance, **b,** sag ratio, **c**, resting potential, **d**, spike latency, **e**, threshold current, **f**, first ISI, **g**, adaptation index, **h**, first spike width, **i**, FI slope across human neurons (*n* = 3 young; *n* = 3 adolescent). **j**-**r** as in (**a**-**i**) but for mouse (*n* = 2 young; *n* = 6 adolescent).

**Extended Data Fig. 5:**
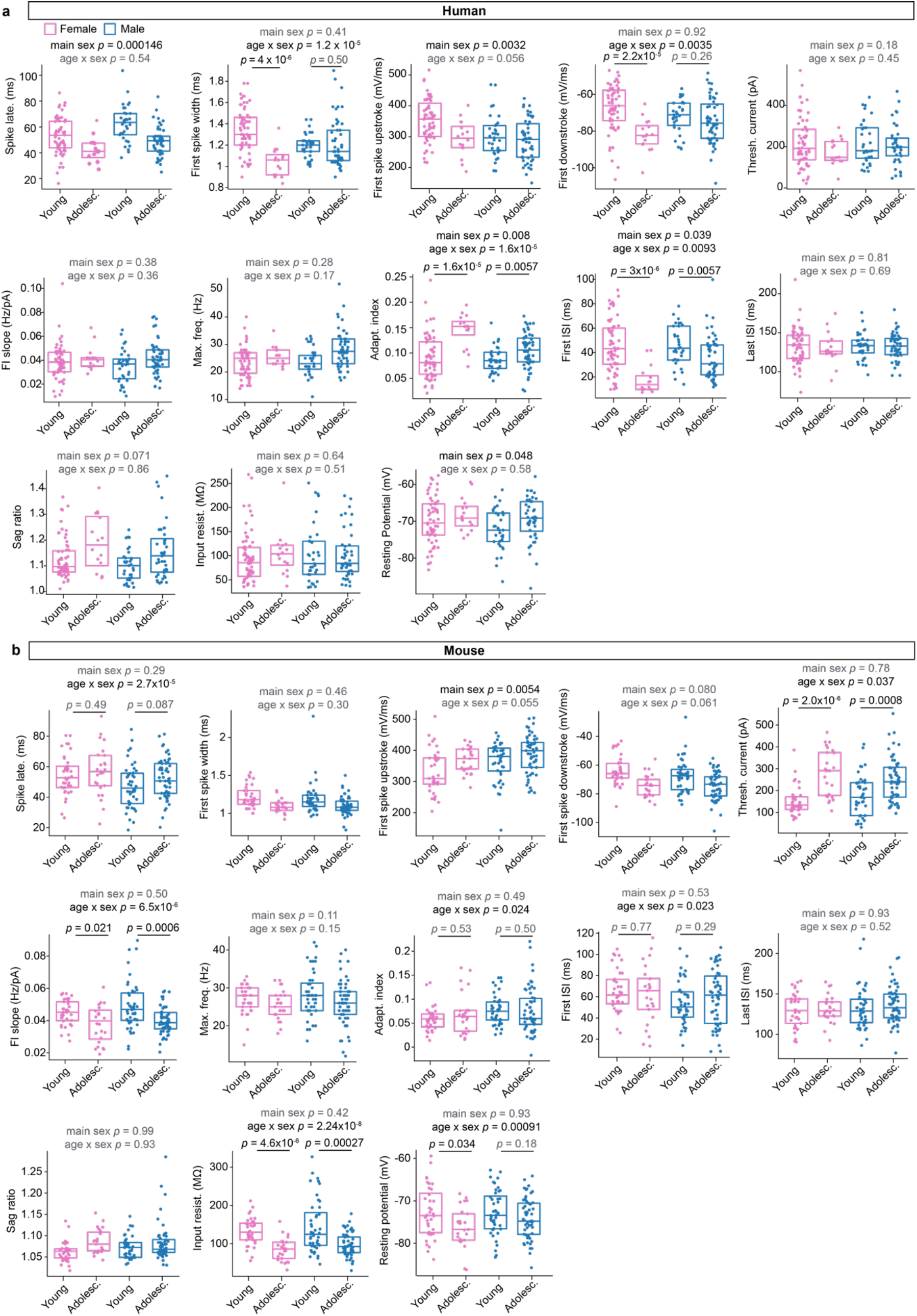
Developmentally-regulated intrinsic physiology properties, disaggregated by sex. Physiology features from Fig. 1 disaggregated by sex in humans (**a**) and mouse (**b**).

**Extended Data Fig. 6:**
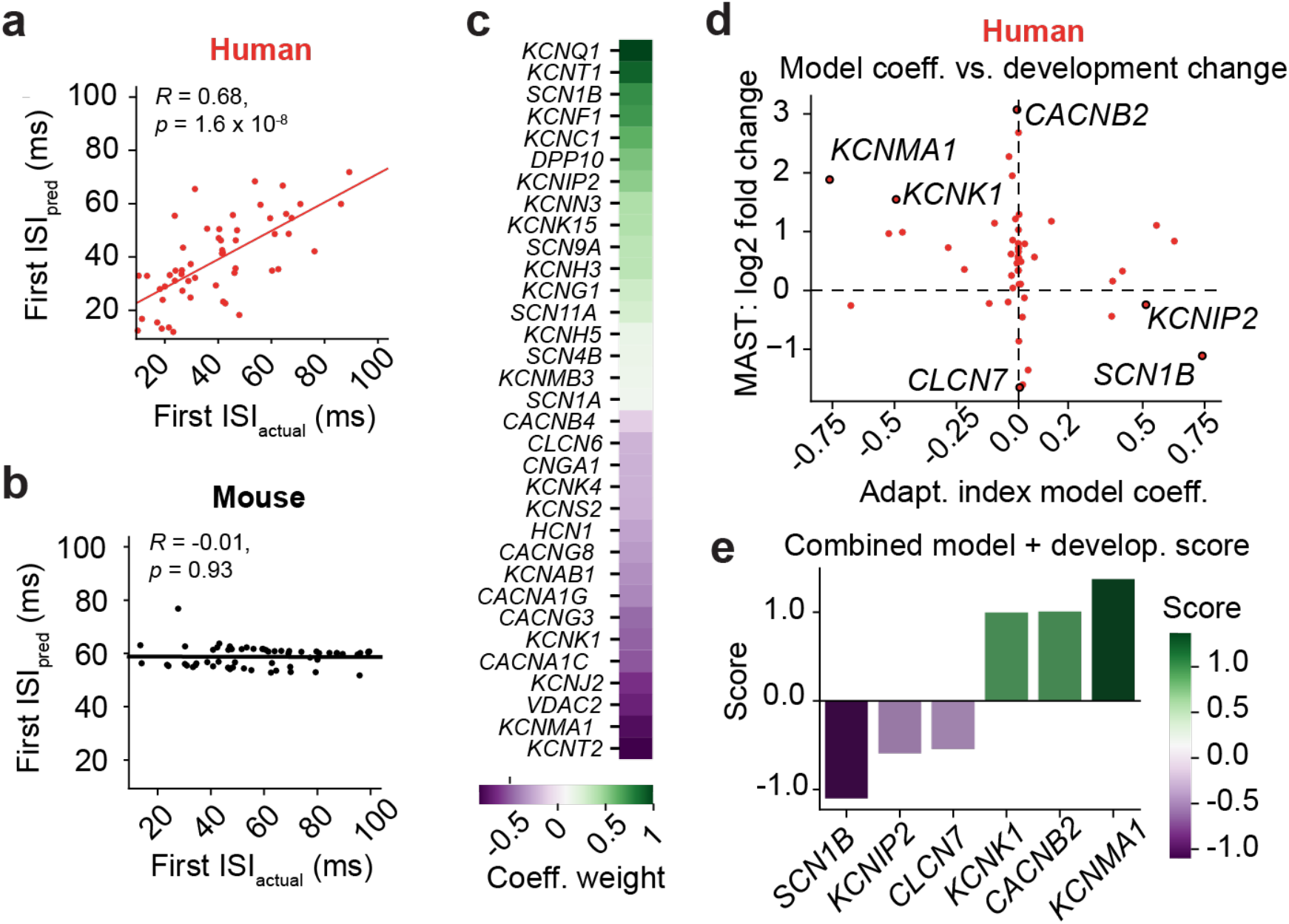
Ion channel gene expression predicts first ISI. Scatter plots denote individual human (*n* = 54) (**a**) or mouse (*n* = 58) (**b**) Patch-Seq neurons, with the *x*-axis indicating the measured first ISI and the *y*-axis the predicted first ISI. Human, but not mouse, model predictions were significantly correlated to the measured first ISI, quantified via Pearson’s *R*. **c**, Non-zero coefficients assigned to genes by the model, with coefficients scaled to the absolute maximum value across all genes. **d**, Scatterplot displays the first ISI linear model coefficient (*x*-axis) versus the development fold change quantified with MAST (*y*-axis) for every ion channel gene input into the human first ISI model. **e**, Sum of development log2fc values and model coefficients scaled to the absolute maximum value.

**Extended Data Fig. 7:**
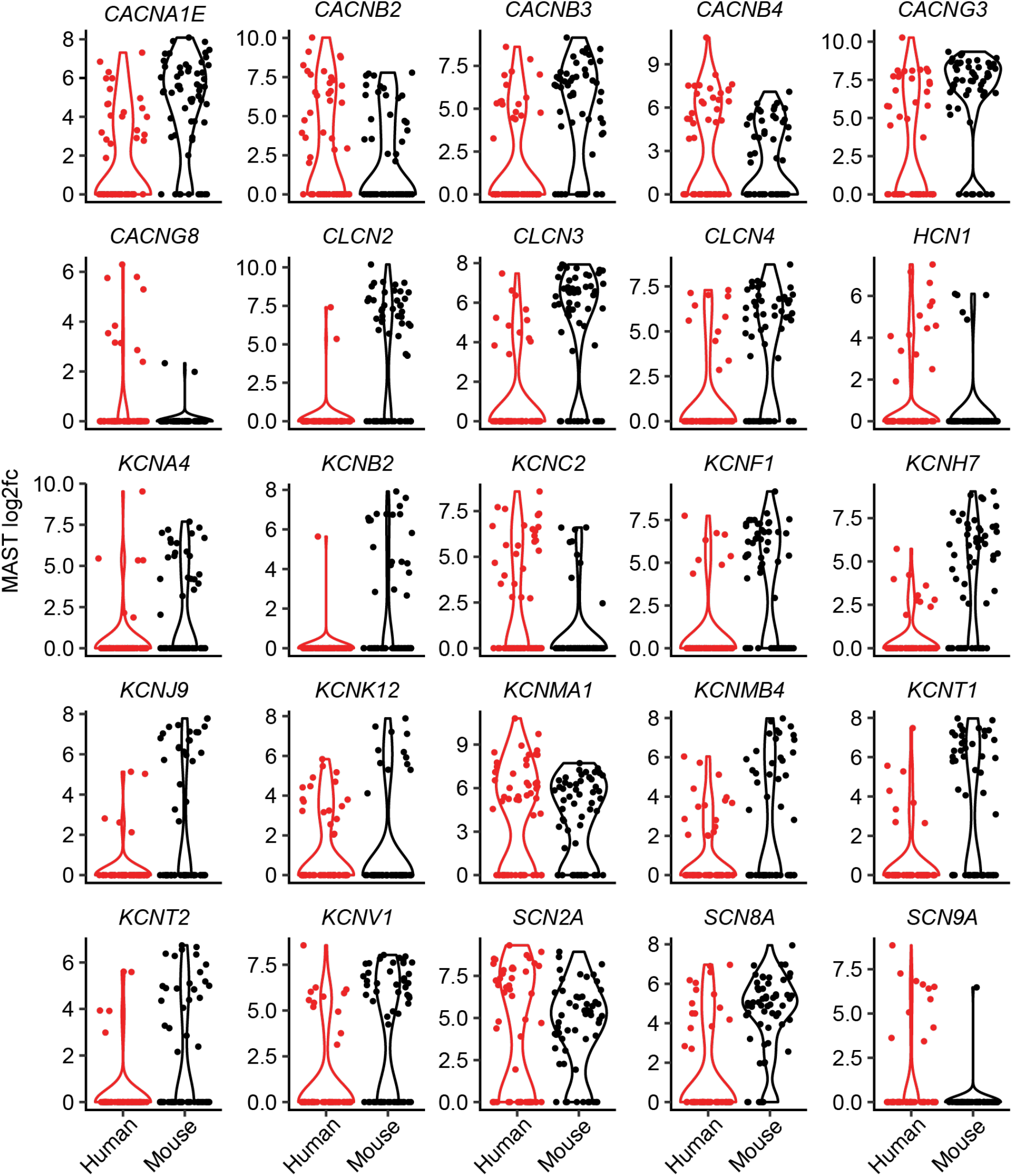
Differentially-expressed ion channel genes across species. Cross-species MAST results for genes that were significantly differentially expressed with *p* < 0.05 after Benjamini & Hochberg False Discovery Rate (FDR) correction.

**Extended Data Fig. 8:**
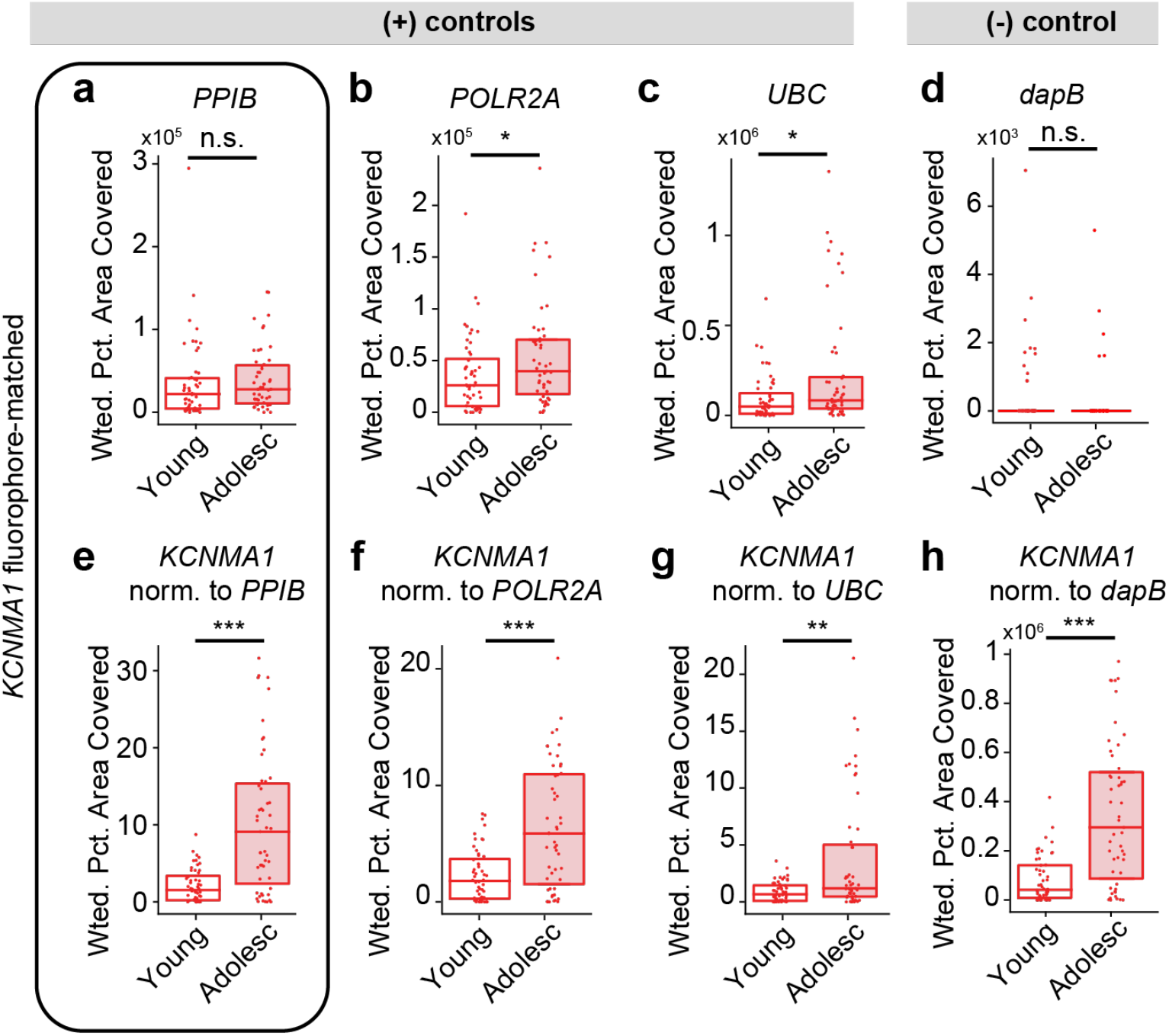
*KCNMA1* upregulation across development cannot be explained by systematic differences in RNA quality or autofluorescence. Developmental expression of three different housekeeping genes: **a**, *PPIB* shows no developmental change (*p* = 0.163); *POLR2A* (**b**) and *UBC* (**c**) show significant upregulation (*p* = 0.043, 0.011, respectively), though the magnitude of the effect is very small. Note that *PPIB* was run in the same channel and using the same fluorophore as *KCNMA1* and thus is the most relevant comparison. **d**, As in (**a**-**c**), but for the negative control bacterial gene, *dapB*. Any signal is presumed autofluorescence or non-specific probe binding. There was no significant change across development in this signal (*p* = 0.12), which is overall very sparse and dim relative to positive housekeeping control genes and *KCNMA1*. **e**, *KCNMA1* expression still increased across development when normalized to the average *PPIB* expression on a per-sample basis (*p* = 1x10^-6^), **f**, As in (**e**) but for *POLR2A*, which still yielded a significant developmental change in *KCNMA1* (*p* = 3.6x10^-5^), **g**, as in (**e**,**f**) but for *UBC*, which still yielded a significant developmental change in *KCNMA1* (*p* = 0.0041), **h**, *KCNMA1* expression normalized to *dapB* negative control expression by subtracting the average signal in (**d**). Subtraction was used instead of division to prevent dividing by zero in some cases with no signal in negative controls. Sample sizes for all experiments were *N* = 4 young human and *N* = 3 adolescent human, with donors evenly and pseudo-randomly subsampled to yield *n* = 50 cells per age. Positive and negative controls were run on samples from the same donors as in Fig. 3, and normalization was applied on a per-donor basis.

**Extended Data Fig. 9:**
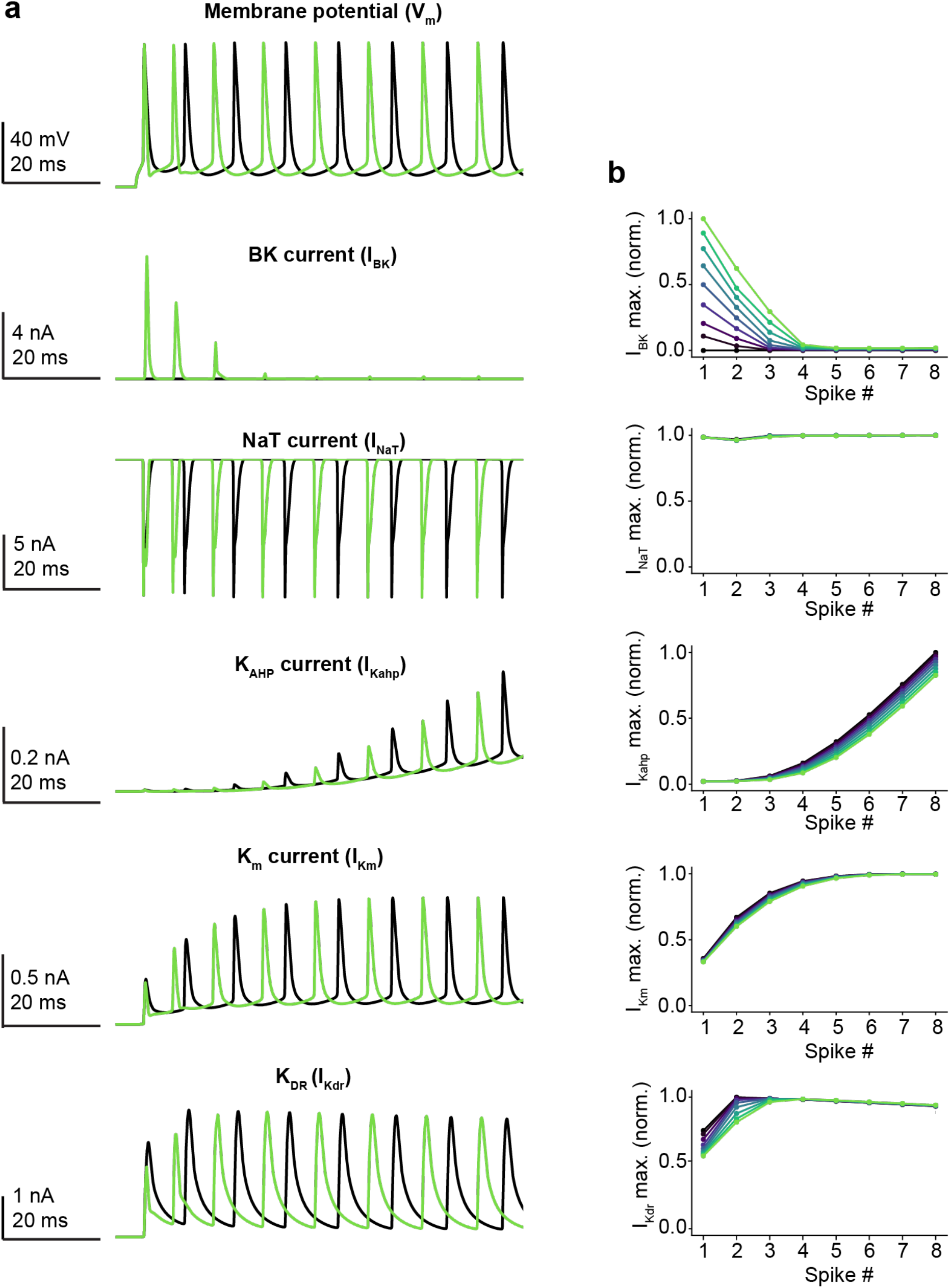
Ion channel currents across the spike train in the multi-compartmental model. a, Membrane potential (top) and current (bottom) for various ion channels involved in spike initiation (INaT) or repolarization (IBK, IKahp, IKm, IKdr). **b**, Maximum value for each current in (**a**) during each spike in the train. INaT was relatively stable across all spikes, further implying that the key feature accelerating initiation of the next spike in the presence of IBK was faster de-inactivation of INaT (Fig. 3), and not overall differences in sodium current levels. Also of note is the reciprocal relationship between IBK and IKdr, which demonstrates that IBK is replaced by the slower IKdr in later spikes in the train.

**Extended Data Fig. 10:**
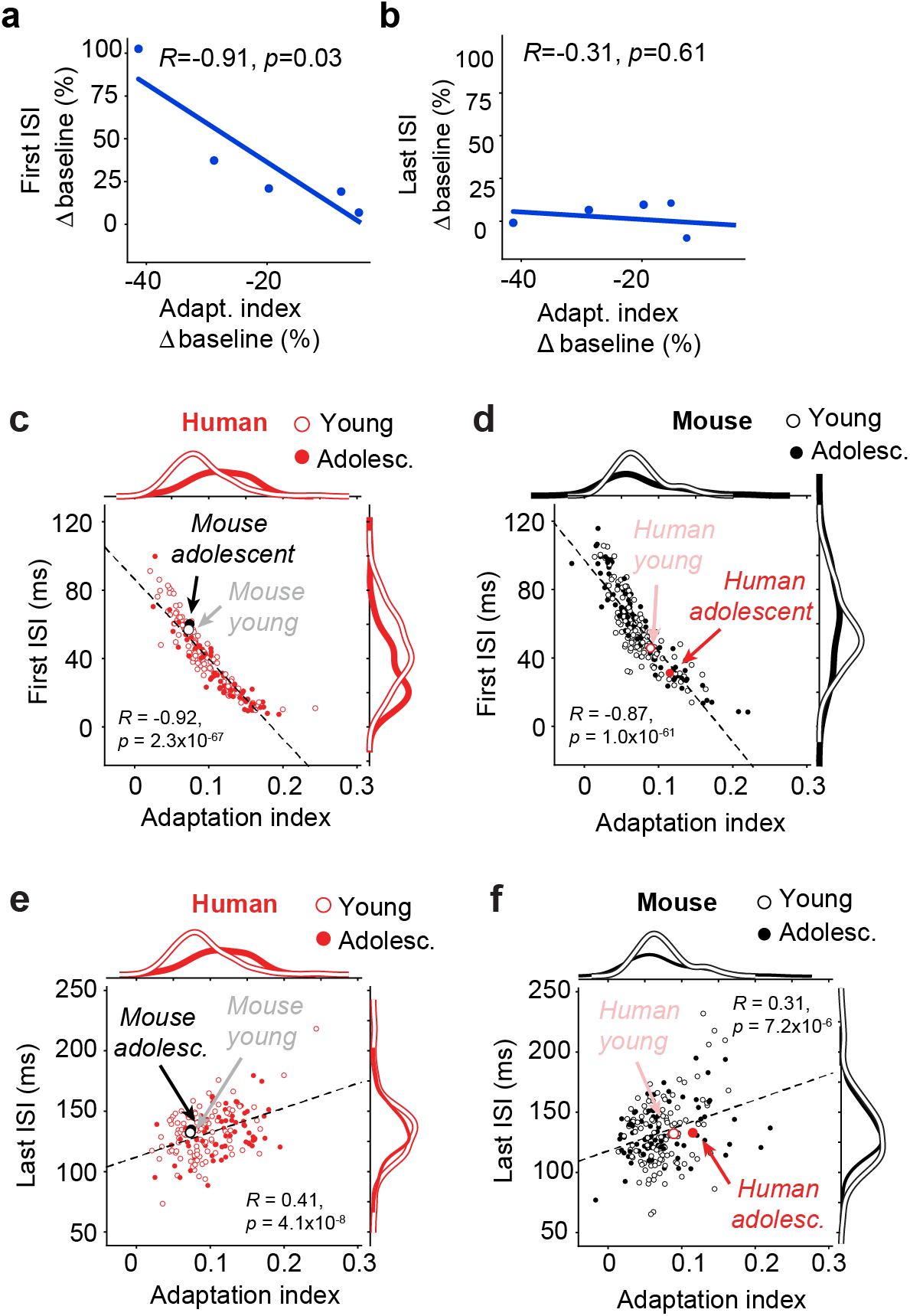
Relationship between first ISI and adaptation after Paxilline blockade and across development and species. a, Percentage-wise change in first ISI versus percentage-wise change in adaptation after Paxilline blockade. **b,** As in (**a**), but for the percentage-wise change in last ISI after Paxilline blockade. The length of the first ISI and the adaptation index are significantly anti-correlated across human (**c**) and mouse (**d**) neurons. This highlights that the relationship between first ISI and adaptation may be conserved across species, but human neurons exclusively showed developmental changes, as well as showed more extreme first ISI and adaptation values relative to mouse, which can be seen by comparing means and distributions. The length of the last ISI and the adaptation index are significantly positively correlated across human (**e**) and mouse (**f**) neurons, albeit with smaller magnitude than (**c**,**d**).

**Extended Data Table 1.** Human neurosurgical resection donor information.

| Age (yrs.) | Sex | Cortical Area | Hemisphere | Diagnosis | Use |
| --- | --- | --- | --- | --- | --- |
| 0.75 | Female | Temporal | L | Epilepsy | Physiology |
| 2 | Male | Temporal | R | Epilepsy | Physiology |
| 2 | Female | Temporal | R | Epilepsy | Physiology |
| 3 | Female | Temporal | L | Epilepsy | Physiology |
| 3 | Female | Temporal | L | Epilepsy | Physiology |
| 4 | Female | Temporal | R | Epilepsy | Physiology |
| 8 | Female | Temporal | L | Epilepsy | Physiology |
| 8 | Female | Temporal | R | Epilepsy | Physiology |
| 8 | Female | Temporal | L | Epilepsy | Physiology |
| 9 | Male | Temporal | R | Epilepsy | Physiology |
| 9 | Male | Temporal | L | Epilepsy | Physiology |
| 9 | Female | Temporal | L | Epilepsy | Physiology |
| 11 | Male | Temporal | L | Epilepsy | Physiology |
| 11 | Female | Temporal | L | Epilepsy | Physiology |
| 13 | Male | Temporal | L | Epilepsy | Physiology |
| 13 | Male | Temporal | R | Epilepsy | Physiology |
| 15 | Male | Temporal | R | Tumor | Physiology |
| 16 | Female | Temporal | L | Epilepsy | Physiology |
| 16 | Male | Temporal | R | Epilepsy | Physiology |
| 16 | Male | Temporal | L | Epilepsy | Physiology+Pharmacology |
| 17 | Male | Temporal | R | Epilepsy | Physiology |
| 17 | Male | Temporal | R | Epilepsy | Physiology |
| 18 | Male | Temporal | R | Epilepsy | Physiology |
| 18 | Male | Temporal | L | Epilepsy | Physiology |
| 1 | Male | Temporal | R | Epilepsy | RNAscope |
| 2 | Male | Temporal | R | Epilepsy | RNAscope |
| 6 | Female | Temporal | L | Epilepsy | RNAscope |
| 8 | Female | Temporal | L | Epilepsy | RNAscope |
| 14 | Male | Temporal | L | Epilepsy | RNAscope |
| 14 | Female | Temporal | R | Epilepsy | RNAscope |
| 14 | Male | Temporal | R | Epilepsy | RNAscope |
| 1 | Male | Frontal | L | Epilepsy | Pharmacology |
| 5 | Female | Frontal | L | Epilepsy | Pharmacology |
| 6 | Male | Frontal | R | Epilepsy | Pharmacology |
| 12 | Male | Temporal | L | Epilepsy | Pharmacology |
| 15 | Male | Frontal | L | Epilepsy | Pharmacology |
| 30 | Male | Temporal | R | Epilepsy | Pharmacology |

**Extended Data Table 2.** Average model coefficients assigned to ion channel genes across train-test splits.

| **Gene** | **Model Coeff. (Avg.)** | **Model Coeff. (Scaled)** |
| --- | --- | --- |
| *CLCN6* | 0.00123718 | 1 |
| *KCNJ2* | 0.00120597 | 0.97477226 |
| *KCNMA1* | 0.00109989 | 0.88902928 |
| *CACNA1C* | 0.00101376 | 0.81941718 |
| *KCNK1* | 0.00088597 | 0.71612077 |
| *CACNG8* | 0.00075277 | 0.60846132 |
| *KCNT2* | 0.00070064 | 0.56632461 |
| *KCNAB1* | 0.00067712 | 0.54731006 |
| *CACNG3* | 0.00063659 | 0.51455019 |
| *VDAC2* | 0.00041355 | 0.33426528 |
| *KCNS2* | 0.00032967 | 0.26646991 |
| *KCNH7* | 0.00016314 | 0.13186348 |
| *KCNC4* | 0.00012824 | 0.10365382 |
| *CACNA2D4* | 0.00011503 | 0.09297463 |
| *KCNJ6* | -0.0001044 | -0.0843949 |
| *KCNK3* | -0.0001675 | -0.1353826 |
| *KCNK15* | -0.0002354 | -0.1902581 |
| *KCNF1* | -0.0002396 | -0.1936464 |
| *SCN9A* | -0.0002402 | -0.1941466 |
| *CACNA1H* | -0.0002712 | -0.2192105 |
| *KCNN3* | -0.0004374 | -0.3535601 |
| *DPP10* | -0.000441 | -0.3564716 |
| *CACNA1A* | -0.000454 | -0.3669717 |
| *KCNAB2* | -0.0004623 | -0.3736475 |
| *SCN2B* | -0.0005219 | -0.4218523 |
| *SCN1A* | -0.0006015 | -0.486169 |
| *KCNMB3* | -0.0007524 | -0.6081563 |
| *KCNIP2* | -0.0007887 | -0.6375283 |
| *SCN1B* | -0.0008379 | -0.6772388 |
| *KCNQ1* | -0.0008897 | -0.719141 |
| *KCNT1* | -0.0009373 | -0.7576497 |
| *KCNC1* | -0.0010327 | -0.8347342 |

## References

1. Rakic, P., et al., Concurrent overproduction of synapses in diverse regions of the primate cerebral cortex. Science, 1986. 232(4747): p. 232–5.

2. Huttenlocher, P.R., Morphometric study of human cerebral cortex development. Neuropsychologia, 1990. 28(6): p. 517–27.

3. Averbeck, B.B., Pruning recurrent neural networks replicates adolescent changes in working memory and reinforcement learning. Proc Natl Acad Sci U S A, 2022. 119(22): p. e2121331119. PMC9295803: PMC9295803

4. Wildenberg, G., et al., Isochronic development of cortical synapses in primates and mice. Nat Commun, 2023. 14(1): p. 8018. PMC10695974: PMC10695974

5. Katz, L.C. and C.J. Shatz, Synaptic activity and the construction of cortical circuits. Science, 1996. 274(5290): p. 1133–8.

6. Cooper, D.L., et al., Synchronized changes to relative neuron populations in postnatal human neocortical development. Cogn Neurodyn, 2010. 4(2): p. 151–63. PMC2866365: PMC2866365

7. Brill, J. and J.R. Huguenard, Sequential changes in AMPA receptor targeting in the developing neocortical excitatory circuit. J Neurosci, 2008. 28(51): p. 13918–28. PMC2706010: PMC2706010

8. Oswald, A.M. and A.D. Reyes, Maturation of intrinsic and synaptic properties of layer 2/3 pyramidal neurons in mouse auditory cortex. J Neurophysiol, 2008. 99(6): p. 2998–3008. PMC3056441: PMC3056441

9. Kalemaki, K., et al., The developmental changes in intrinsic and synaptic properties of prefrontal neurons enhance local network activity from the second to the third postnatal weeks in mice. Cereb Cortex, 2022. 32(17): p. 3633–3650.

10. Kang, J., J.R. Huguenard, and D.A. Prince, Development of BK channels in neocortical pyramidal neurons. J Neurophysiol, 1996. 76(1): p. 188–98.

11. Huguenard, J.R., O.P. Hamill, and D.A. Prince, Developmental changes in Na+ conductances in rat neocortical neurons: appearance of a slowly inactivating component. J Neurophysiol, 1988. 59(3): p. 778–95.

12. Wang, L., et al., A cross-species proteomic map reveals neoteny of human synapse development. Nature, 2023. 622(7981): p. 112–119. PMC10576238: PMC10576238

13. Petanjek, Z., et al., Extraordinary neoteny of synaptic spines in the human prefrontal cortex. Proc Natl Acad Sci U S A, 2011. 108(32): p. 13281–6. PMC3156171: PMC3156171

14. Vermaercke, B., et al., SYNGAP1 deficiency disrupts synaptic neoteny in xenotransplanted human cortical neurons in vivo. Neuron, 2024. 112(18): p. 3058–3068 e8. PMC11446607: PMC11446607

15. Vanderhaeghen, P. and F. Polleux, Developmental mechanisms underlying the evolution of human cortical circuits. Nat Rev Neurosci, 2023. 24(4): p. 213–232. PMC10064077: PMC10064077

16. Wang, L., et al., Molecular and cellular dynamics of the developing human neocortex. Nature, 2025. 647(8088): p. 169–178. PMC12589127 Therapeutics. J.L. is a co-founder and a member of the scientific advisory board of SensOmics, Inc. The other authors declare no competing interests.: PMC12589127 Therapeutics. J.L. is a co-founder and a member of the scientific advisory board of SensOmics, Inc. The other authors declare no competing interests.

17. Libe-Philippot, B., et al., Synaptic neoteny of human cortical neurons requires species-specific balancing of SRGAP2-SYNGAP1 cross-inhibition. Neuron, 2024. 112(21): p. 3602–3617 e9. PMC11546603: PMC11546603

18. Charrier, C., et al., Inhibition of SRGAP2 function by its human-specific paralogs induces neoteny during spine maturation. Cell, 2012. 149(4): p. 923–35. PMC3357949: PMC3357949

19. Velmeshev, D., et al., Single-cell analysis of prenatal and postnatal human cortical development. Science, 2023. 382(6667): p. eadf0834. PMC11005279: PMC11005279

20. Schwarz, L.A., et al., Cortical development dynamics across autism spectrum disorder mouse models. Nature, 2026.

21. Antoine, M.W., et al., Increased Excitation-Inhibition Ratio Stabilizes Synapse and Circuit Excitability in Four Autism Mouse Models. Neuron, 2019. 101(4): p. 648–661 e4. PMC6733271: PMC6733271

22. Greenberg, B.D., et al., Altered cortical excitability in obsessive-compulsive disorder. Neurology, 2000. 54(1): p. 142–7.

23. Shmelkov, S.V., et al., Slitrk5 deficiency impairs corticostriatal circuitry and leads to obsessive-compulsive-like behaviors in mice. Nat Med, 2010. 16(5): p. 598–602, 1p following 602. PMC2907076: PMC2907076

24. Ahmari, S.E., et al., Repeated cortico-striatal stimulation generates persistent OCD-like behavior. Science, 2013. 340(6137): p. 1234–9. PMC3954809: PMC3954809

25. Hamm, J.P., et al., Altered Cortical Ensembles in Mouse Models of Schizophrenia. Neuron, 2017. 94(1): p. 153–167 e8. PMC5394986: PMC5394986

26. Kushner, J.K., et al., Characterizing the Diversity of Layer 2/3 Human Neocortical Neurons in Pediatric Epilepsy. eNeuro, 2025. 12(5). PMC12061357: PMC12061357

27. Verhoog, M.B., et al., Human neurons undergo protracted functional maturation into adulthood. bioRxiv, 2025.

28. Barzo, P., et al., Electrophysiology and morphology of human cortical supragranular pyramidal cells in a wide age range. Elife, 2025. 13. PMC11952751: PMC11952751

29. Cadwell, C.R., et al., Electrophysiological, transcriptomic and morphologic profiling of single neurons using Patch-seq. Nat Biotechnol, 2016. 34(2): p. 199–203. PMC4840019: PMC4840019

30. Foldy, C., et al., Single-cell RNAseq reveals cell adhesion molecule profiles in electrophysiologically defined neurons. Proc Natl Acad Sci U S A, 2016. 113(35): p. E5222–31. PMC5024636: PMC5024636

31. Fuzik, J., et al., Integration of electrophysiological recordings with single-cell RNA-seq data identifies neuronal subtypes. Nat Biotechnol, 2016. 34(2): p. 175–183. PMC4745137: PMC4745137

32. van Loo, K.M.J., et al., What makes the human brain special: from cellular function to clinical translation. J Neurophysiol, 2025. 134(4): p. 1197–1213.

33. Berg, J., et al., Human neocortical expansion involves glutamatergic neuron diversification. Nature, 2021. 598(7879): p. 151–158. PMC8494638: PMC8494638

34. Szegedi, V., et al., HCN channels at the cell soma ensure the rapid electrical reactivity of fast-spiking interneurons in human neocortex. PLoS Biol, 2023. 21(2): p. e3002001. PMC9934405: PMC9934405

35. Goriounova, N.A., et al., Large and fast human pyramidal neurons associate with intelligence. Elife, 2018. 7. PMC6363383: PMC6363383

36. Kalmbach, B.E., et al., *h-Channels Contribute to Divergent Intrinsic Membrane Properties of Supragranular Pyramidal Neurons in Human versus Mouse Cerebral Cortex*. Neuron, 2018. 100(5): p. 1194–1208 e5. PMC6447369: PMC6447369

37. Testa-Silva, G., et al., High bandwidth synaptic communication and frequency tracking in human neocortex. PLoS Biol, 2014. 12(11): p. e1002007. PMC4244038: PMC4244038

38. Campagnola, L., et al., Local connectivity and synaptic dynamics in mouse and human neocortex. Science, 2022. 375(6585): p. eabj5861. PMC9970277: PMC9970277

39. Peng, Z., et al., Directed acyclic graph for epidemiological studies in childhood food allergy: Construction, user’s guide, and application. Allergy, 2024. 79(8): p. 2051–2064.

40. Eyal, G., et al., Unique membrane properties and enhanced signal processing in human neocortical neurons. Elife, 2016. 5. PMC5100995: PMC5100995

41. Deitcher, Y., et al., Comprehensive Morpho-Electrotonic Analysis Shows 2 Distinct Classes of L2 and L3 Pyramidal Neurons in Human Temporal Cortex. Cereb Cortex, 2017. 27(11): p. 5398–5414. PMC5939232: PMC5939232

42. McCormick, D.A., et al., Comparative electrophysiology of pyramidal and sparsely spiny stellate neurons of the neocortex. J Neurophysiol, 1985. 54(4): p. 782–806.

43. Glasser, M.F., et al., Trends and properties of human cerebral cortex: correlations with cortical myelin content. Neuroimage, 2014. 93 **Pt** **2**: p. 165–75. PMC3795824: PMC3795824

44. Glasser, M.F., et al., A multi-modal parcellation of human cerebral cortex. Nature, 2016. 536(7615): p. 171–178. PMC4990127: PMC4990127

45. Planert, H., et al., Electrophysiological classification of human layer 2-3 pyramidal neurons reveals subtype-specific synaptic interactions. Nat Neurosci, 2026. 29(2): p. 455–466. PMC12880919: PMC12880919

46. Bomkamp, C., et al., Transcriptomic correlates of electrophysiological and morphological diversity within and across excitatory and inhibitory neuron classes. PLoS Comput Biol, 2019. 15(6): p. e1007113. PMC6599125: PMC6599125

47. Tripathy, S.J., et al., Transcriptomic correlates of neuron electrophysiological diversity. PLoS Comput Biol, 2017. 13(10): p. e1005814. PMC5673240: PMC5673240

48. Gouwens, N.W., et al., Integrated Morphoelectric and Transcriptomic Classification of Cortical GABAergic Cells. Cell, 2020. 183(4): p. 935–953 e19. PMC7781065: PMC7781065

49. Gu, N., K. Vervaeke, and J.F. Storm, BK potassium channels facilitate high-frequency firing and cause early spike frequency adaptation in rat CA1 hippocampal pyramidal cells. J Physiol, 2007. 580(Pt.3): p. 859–82. PMC2075463: PMC2075463

50. Wang, B., et al., A Subtype of Inhibitory Interneuron with Intrinsic Persistent Activity in Human and Monkey Neocortex. Cell Rep, 2015. 10(9): p. 1450–1458.

51. Mohan, H., et al., Dendritic and Axonal Architecture of Individual Pyramidal Neurons across Layers of Adult Human Neocortex. Cereb Cortex, 2015. 25(12): p. 4839–53. PMC4635923: PMC4635923

52. Kail, R., Developmental change in speed of processing during childhood and adolescence. Psychol Bull, 1991. 109(3): p. 490–501.

53. Kail, R.V. and E. Ferrer, Processing speed in childhood and adolescence: longitudinal models for examining developmental change. Child Dev, 2007. 78(6): p. 1760–70.

54. Gonzalez-Burgos, G., et al., Distinct Properties of Layer 3 Pyramidal Neurons from Prefrontal and Parietal Areas of the Monkey Neocortex. J Neurosci, 2019. 39(37): p. 7277–7290. PMC6759021: PMC6759021

55. Mease, R.A., et al., Emergence of adaptive computation by single neurons in the developing cortex. J Neurosci, 2013. 33(30): p. 12154–70. PMC3721832: PMC3721832

56. Zhang, H., et al., The noise cancelation effects caused by spike-frequency adaptation in single neurons. Nonlinear Dynamics, 2020. 100(2): p. 1825–1835.

57. Higgs, M.H., S.J. Slee, and W.J. Spain, Diversity of gain modulation by noise in neocortical neurons: regulation by the slow afterhyperpolarization conductance. J Neurosci, 2006. 26(34): p. 8787–99. PMC6674385: PMC6674385

58. Benda, J. and R.M. Hennig, Spike-frequency adaptation generates intensity invariance in a primary auditory interneuron. J Comput Neurosci, 2008. 24(2): p. 113–36.

59. Paraskevopoulou, F., et al., Impaired inhibitory GABAergic synaptic transmission and transcription studied in single neurons by Patch-seq in Huntington’s disease. Proc Natl Acad Sci U S A, 2021. 118(19). PMC8126788: PMC8126788

60. Scala, F., et al., Phenotypic variation of transcriptomic cell types in mouse motor cortex. Nature, 2021. 598(7879): p. 144-150. PMC8113357: PMC8113357

61. Golowasch, J., et al., Failure of averaging in the construction of a conductance-based neuron model. J Neurophysiol, 2002. 87(2): p. 1129–31.

62. Schulz, D.J., J.M. Goaillard, and E.E. Marder, Quantitative expression profiling of identified neurons reveals cell-specific constraints on highly variable levels of gene expression. Proc Natl Acad Sci U S A, 2007. 104(32): p. 13187–91. PMC1933263: PMC1933263

63. Dueck, H., et al., Deep sequencing reveals cell-type-specific patterns of single-cell transcriptome variation. Genome Biol, 2015. 16(1): p. 122. PMC4480509: PMC4480509

64. Cheng, S., et al., Vision-dependent specification of cell types and function in the developing cortex. Cell, 2022. 185(2): p. 311–327 e24. PMC8813006: PMC8813006

65. Yao, Z., et al., A taxonomy of transcriptomic cell types across the isocortex and hippocampal formation. Cell, 2021. 184(12): p. 3222–3241 e26. PMC8195859: PMC8195859

66. Ramsden, H.L., et al., Laminar and dorsoventral molecular organization of the medial entorhinal cortex revealed by large-scale anatomical analysis of gene expression. PLoS Comput Biol, 2015. 11(1): p. e1004032. PMC4304787: PMC4304787

67. O’Leary, T.P., et al., Extensive and spatially variable within-cell-type heterogeneity across the basolateral amygdala. Elife, 2020. 9. PMC7486123: PMC7486123

68. Cembrowski, M.S., et al., Spatial Gene-Expression Gradients Underlie Prominent Heterogeneity of CA1 Pyramidal Neurons. Neuron, 2016. 89(2): p. 351–68.

69. Kendrick, R.M., S.W. Linderman, and S. Owen, Transcriptomically-measured gene expression predicts physiological variation across single neurons in humans and mice. bioRxiv, 2024.

70. Tibshirani, R., Regression shrinkage and selection via the Lasso. Journal of the Royal Statistical Society Series B-Statistical Methodology, 1996. 58(1): p. 267–288.

71. Finak, G., et al., MAST: a flexible statistical framework for assessing transcriptional changes and characterizing heterogeneity in single-cell RNA sequencing data. Genome Biol, 2015. 16: p. 278. PMC4676162: PMC4676162

72. Plante, A.E., J.P. Whitt, and A.L. Meredith, BK channel activation by L-type Ca(2+) channels Ca(V)1.2 and Ca(V)1.3 during the subthreshold phase of an action potential. J Neurophysiol, 2021. 126(2): p. 427–439. PMC8409951: PMC8409951

73. Vivas, O., et al., Proximal clustering between BK and Ca(V)1.3 channels promotes functional coupling and BK channel activation at low voltage. Elife, 2017. 6. PMC5503510: PMC5503510

74. Li, B., et al., Neuronal Inactivity Co-opts LTP Machinery to Drive Potassium Channel Splicing and Homeostatic Spike Widening. Cell, 2020. 181(7): p. 1547–1565 e15. PMC9310388: PMC9310388

75. Chen, C., et al., Mice lacking sodium channel beta1 subunits display defects in neuronal excitability, sodium channel expression, and nodal architecture. J Neurosci, 2004. 24(16): p. 4030–42. PMC6729427: PMC6729427

76. Baroni, D., et al., Antisense-mediated post-transcriptional silencing of SCN1B gene modulates sodium channel functional expression. Biol Cell, 2014. 106(1): p. 13–29.

77. Deng, P.Y., et al., FMRP regulates neurotransmitter release and synaptic information transmission by modulating action potential duration via BK channels. Neuron, 2013. 77(4): p. 696–711. PMC3584349: PMC3584349

78. Gray, D.A. and J. Woulfe, Lipofuscin and aging: a matter of toxic waste. Sci Aging Knowledge Environ, 2005. 2005(5): p. re1.

79. Bader, P.L., et al., Mouse model of Timothy syndrome recapitulates triad of autistic traits. Proc Natl Acad Sci U S A, 2011. 108(37): p. 15432–7. PMC3174658: PMC3174658

80. Splawski, I., et al., Ca(V)1.2 calcium channel dysfunction causes a multisystem disorder including arrhythmia and autism. Cell, 2004. 119(1): p. 19–31.

81. Chen, X., et al., Antisense oligonucleotide therapeutic approach for Timothy syndrome. Nature, 2024. 628(8009): p. 818–825. PMC11043036 organoids/spheroids and assembloids (listing S.P.P., F.B. as inventors), a patent application for ASO (listing S.P.P., X.C. and F.B. as inventors) and a patent application for transplantation of organoids (listing S.P.P. and O.R. as inventors). PMC11043036 organoids/spheroids and assembloids (listing S.P.P., F.B. as inventors), a patent application for ASO (listing S.P.P., X.C. and F.B. as inventors) and a patent application for transplantation of organoids (listing S.P.P. and O.R. as inventors).

82. Wu, H., et al., Phenotype-to-genotype approach reveals head-circumference-associated genes in an autism spectrum disorder cohort. Clin Genet, 2020. 97(2): p. 338–346. PMC7307605: PMC7307605

83. Laumonnier, F., et al., Association of a functional deficit of the BKCa channel, a synaptic regulator of neuronal excitability, with autism and mental retardation. Am J Psychiatry, 2006. 163(9): p. 1622–9.

84. Song, T., et al., CRL4 antagonizes SCFFbxo7-mediated turnover of cereblon and BK channel to regulate learning and memory. PLoS Genet, 2018. 14(1): p. e1007165. PMC5800687: PMC5800687

85. Deng, P.Y. and V.A. Klyachko, Genetic upregulation of BK channel activity normalizes multiple synaptic and circuit defects in a mouse model of fragile X syndrome. J Physiol, 2016. 594(1): p. 83–97. PMC4704506: PMC4704506

86. Myrick, L.K., et al., Independent role for presynaptic FMRP revealed by an FMR1 missense mutation associated with intellectual disability and seizures. Proc Natl Acad Sci U S A, 2015. 112(4): p. 949–56. PMC4313821: PMC4313821

87. Hebert, B., et al., Rescue of fragile X syndrome phenotypes in Fmr1 KO mice by a BKCa channel opener molecule. Orphanet J Rare Dis, 2014. 9: p. 124. PMC4237919: PMC4237919

88. Libe-Philippot, B., et al., LRRC37B is a human modifier of voltage-gated sodium channels and axon excitability in cortical neurons. Cell, 2023. 186(26): p. 5766–5783 e25. PMC10754148: PMC10754148

89. Williams, S.R. and G.J. Stuart, Site independence of EPSP time course is mediated by dendritic I(h) in neocortical pyramidal neurons. J Neurophysiol, 2000. 83(5): p. 3177–82.

90. Magee, J.C., Dendritic hyperpolarization-activated currents modify the integrative properties of hippocampal CA1 pyramidal neurons. J Neurosci, 1998. 18(19): p. 7613–24. PMC6793032: PMC6793032

91. Dembrow, N.C., B.V. Zemelman, and D. Johnston, Temporal dynamics of L5 dendrites in medial prefrontal cortex regulate integration versus coincidence detection of afferent inputs. J Neurosci, 2015. 35(11): p. 4501–14. PMC4363381: PMC4363381

92. Vaidya, S.P. and D. Johnston, Temporal synchrony and gamma-to-theta power conversion in the dendrites of CA1 pyramidal neurons. Nat Neurosci, 2013. 16(12): p. 1812–20. PMC3958963: PMC3958963

93. Benda, J., A. Longtin, and L. Maler, Spike-frequency adaptation separates transient communication signals from background oscillations. J Neurosci, 2005. 25(9): p. 2312–21. PMC6726095: PMC6726095

94. Prescott, S.A. and T.J. Sejnowski, Spike-rate coding and spike-time coding are affected oppositely by different adaptation mechanisms. J Neurosci, 2008. 28(50): p. 13649–61. PMC2819463: PMC2819463

95. Prescott, S.A., et al., Nonlinear interaction between shunting and adaptation controls a switch between integration and coincidence detection in pyramidal neurons. J Neurosci, 2006. 26(36): p. 9084–97. PMC2913017: PMC2913017

96. Puccini, G.D., M.V. Sanchez-Vives, and A. Compte, Selective detection of abrupt input changes by integration of spike-frequency adaptation and synaptic depression in a computational network model. J Physiol Paris, 2006. 100(1-3): p. 1–15.

97. Panzeri, S., et al., Speed of feedforward and recurrent processing in multilayer networks of integrate-and-fire neurons. Network, 2001. 12(4): p. 423–40.

98. Salaj, D., et al., Spike frequency adaptation supports network computations on temporally dispersed information. Elife, 2021. 10. PMC8313230: PMC8313230

99. Oshlack, A. and M.J. Wakefield, Transcript length bias in RNA-seq data confounds systems biology. Biol Direct, 2009. 4: p. 14. PMC2678084: PMC2678084

100. Bakos, E., et al., Adaptations of the axon initial segment in fast-spiking interneurons of the human neocortex support low action potential thresholds. PLoS Biol, 2025. 23(12): p. e3003549. PMC12798858: PMC12798858

101. Kalmbach, B.E., et al., Signature morpho-electric, transcriptomic, and dendritic properties of human layer 5 neocortical pyramidal neurons. Neuron, 2021. 109(18): p. 2914–2927 e5. PMC8570452: PMC8570452

102. Gomez-de-Mariscal, E., et al., Use of the p-values as a size-dependent function to address practical differences when analyzing large datasets. Sci Rep, 2021. 11(1): p. 20942. PMC8536742: PMC8536742

103. Ancaten-Gonzalez, C., et al., BK channels mediate a presynaptic form of mGluR-LTD in the neonatal hippocampus. Proc Natl Acad Sci U S A, 2025. 122(2): p. e2411506122. PMC11745352: PMC11745352

104. Hines, M.L. and N.T. Carnevale, The NEURON simulation environment. Neural Comput, 1997. 9(6): p. 1179–209.

